# A dopamine gradient controls access to distributed working memory in monkey cortex

**DOI:** 10.1101/2020.09.07.286500

**Authors:** Sean Froudist-Walsh, Daniel P. Bliss, Xingyu Ding, Lucija Jankovic-Rapan, Meiqi Niu, Kenneth Knoblauch, Karl Zilles, Henry Kennedy, Nicola Palomero-Gallagher, Xiao-Jing Wang

## Abstract

Dopamine is critical for working memory. However, its effects throughout the large-scale primate cortex are poorly understood. Here we report that dopamine receptor density per neuron, measured by receptor autoradiography in the macaque monkey cortex, displays a macroscopic gradient along the cortical hierarchy. We developed a connectome- and biophysically-based model for distributed working memory that incorporates multiple neuron types and a dopamine gradient. The model captures an inverted U-shaped dependence of working memory on dopamine. The spatial distribution of mnemonic persistent activity matches that observed in over 90 experimental studies. We show that dopamine filters out irrelevant stimuli by enhancing inhibition of pyramidal cell dendrites. The level of cortical dopamine can also determine whether memory encoding is through persistent activity or an internal synaptic state. Taken together, our work represents a cross-level understanding that links molecules, cell types, recurrent circuit dynamics and a core cognitive function distributed across the cortex.

## Introduction

Our ability to think through difficult problems without distraction is a hallmark of cognition. When faced with a constant stream of information, we must keep certain information in mind and protect it from distraction. For instance, while following a conversation, it is important to focus on and remember the words that you are listening to, while ignoring other sights and sounds around you. This brain function is called working memory. The underlying neural representation engages information-specific persistent neural activity which is internally sustained in the absence of external stimulation across multiple cortical and subcortical areas (Christophel et al. 2017; Courtney et al. 1997; Dotson et al. 2018; Funahashi et al. 1989; Fuster 1973; Guo et al. 2017; Kamiński and Rutishauser 2020; Konecky et al. 2017; Leavitt et al. 2017; Romo et al. 1999; Wang 2001; Warden and Miller 2010).

Working memory and the prefrontal cortex are under the influence of monoaminergic modulation (Robbins and Arnsten 2009). In fact, depletion of dopamine from the prefrontal cortex causes working memory deficits as severe as those seen by lesioning the prefrontal cortex (Brozoski et al. 1979). Dopamine neurons fire in response to stimuli that predict reward, but do not fire persistently during the delay period of working memory tasks (Cohen et al. 2012; Schultz et al. 1993). How might dopamine affect working memory if dopaminergic neurons do not fire during the delay? After dopamine neurons fire, dopamine is released, and dopamine levels remain elevated in the cortex for hundreds of milliseconds to tens of seconds, before being slowly cleared away (Cass and Gerhardt 1995; Garris and Wightman 1994; Muller et al. 2014; Mundorf et al. 2001). Prefrontal neuron activity during working memory depends on precise levels of activation of the dopamine D1 receptors, with both too little and too much D1 receptor stimulation disrupting delay period activity (Vijayraghavan et al. 2007; Wang et al. 2019; Williams and Goldman-Rakic 1995).

Experimental and modelling studies of dopamine on persistent activity in working memory have focused on isolated local brain regions, generally in the lateral prefrontal cortex (Brunel and Wang 2001; Durstewitz et al. 2000; Jacob et al. 2016; Vijayraghavan et al. 2016, 2007; Wang et al. 2019; Williams and Goldman-Rakic 1995), where it has been shown that dopamine can enhance persistent activity through its effects on the NMDA and GABA receptors (Seamans et al. 2001a,b; Seamans and Yang 2004; Wang et al. 2013). If the effects of dopamine are truly restricted to small areas of cortex, then dopamine seems unlikely to be able to engage the widespread distributed activity seen during active working memory. If dopamine could enable persistent activity across large parts of cortex, then firing of dopamine neurons in response to behaviourally relevant stimuli could be a candidate mechanism to engage active working memory.

In spite of progress, our understanding remains far from complete. In this work, we tackled two open questions. First, how does dopamine modulate working memory across a multi-regional large-scale cortical system? Dopamine modulates neural activity through its receptors, of which the D1 receptor is the most common in cortex. The density of D1 receptors is known only for small sections of monkey cortex (Goldman-Rakic et al. 1990; Impieri et al. 2019; Lidow et al. 1991; Niu et al. 2020; Richfield et al. 1989). A detailed map of the density of D1 receptors across cortex would clarify the degree to which dopamine’s influence on cortical processing is restricted to specific sections of cortex, or distributed throughout cortical systems.

Second, does dopamine contribute to robust working memory against distractors by virtue of differential impacts on different cell types? Drugs that promote synthesis or prevent reuptake of dopamine at the synapse make working memory less vulnerable to distracting stimuli (Fallon et al. 2017, 2016), and D1-agonists and antagonists determine how strongly neurons in prefrontal cortex encode distractor stimuli (Jacob et al. 2016). An early theoretical study proposed that inhibition targeted more strongly towards the dendrites, and away from the soma of pyramidal cells could increase the resistance of working memory to distraction (Wang et al. 2004a). Distinct inhibitory cell types primarily focus their inhibition on the dendrites or soma of pyramidal cells, or on other inhibitory neurons (Adesnik et al. 2012; Jiang et al. 2015; Pfeffer et al. 2013; Tremblay et al. 2016). One intriguing possibility is that dopamine may shift the balance between distinct types of inhibitory neurons in order to keep distracting information from working memory.

We first set out to examine whether dopamine D1 receptor densities across cortex represent random heterogeneity, or a systematic gradient of receptor expression. Using in-vitro autoradiography, we measured the density of dopamine D1 receptors across 109 areas of macaque cortex. We found a systematic gradient of D1 receptors, that increased along the cortical hierarchy and peaked in frontal and parietal cortices. We then built a large-scale computational model of macaque cortex that is endowed with multiple cell types and is capable of performing working memory tasks. Interactions between multiple cell types across cortex were modulated by the density of dopamine D1 receptor receptors, and constrained by retrograde tract-tracing data. We investigated whether dopamine could ignite active working memory representations across frontal and parietal cortex given the measured pattern of D1 receptor receptors and cortico-cortical connections. We found that sufficient dopamine release in the model led to persistent working memory activity throughout the cortex that closely matched experimentally observed mnemonic activity in nearly 20 cortical areas in macaque monkeys. Stimuli that evoked too little or too much dopamine release could still be remembered, but stored in an internal synaptic state instead of persistent neural firing, consistent with an activity-silent scenario (Rose et al. 2016; Stokes 2015; Wolff et al. 2017). In this case, however, memories were more vulnerable to distractions. Appropriate dopamine release in response to relevant and distracting stimuli could be learned through reinforcement. Our model simulations suggest that dopamine may render memories robust to distraction by altering interactions between distinct types of inhibitory cells, blocking external stimuli from entering frontal cortex and promoting recurrent excitatory activity. Together, our results uncover a macroscopic gradient of dopamine D1 receptor distribution and elucidate how differential dopamine actions on different cell types ensure distractor-resistant working memory activity throughout primate cortex.

## Results

### A hierarchical gradient of dopamine D1 receptors per neuron across monkey cortex

To investigate how dopamine modulation may vary across cortical regions, we first analized D1 and D2 receptor distribution patterns throughout the macaque brain using in-vitro receptor autoradiography (Fig. S1; Supplementary Data). Autoradiography enables the quantification of endogenous receptors in the cell membrane through the use of radioactive ligands (Palomero-Gallagher and Zilles 2018). The highest densities (in fmol/mg protein) of both receptor types were found in the basal ganglia, with the caudate nucleus (D1 298 ± 28; D2 188 ± 30) and putamen (D1 273 ± 40; D2 203 ± 37) presenting considerably higher values than the internal (D1 97 ± 34; D2 22 ± 12) or external (D1 55 ± 16; D2 30 ± 11) subdivisions of the globus pallidus. Cortical D1 receptor densities ranged from 49 ± 13 fmol/mg protein in area 4a of the primary motor cortex to 101 ± 35 fmol/mg protein in orbitofrontal area 11l. We found a systematic gradient of D1 receptors across the 109 cytoarchitectonically identified cortical areas, whereby densities reached maximal values in areas of the frontal and parietal cortex (Fig. 1A). The density of the D2 receptor in cortex is so low that it is not detectable by means of the here applied method.

**Figure 1:**
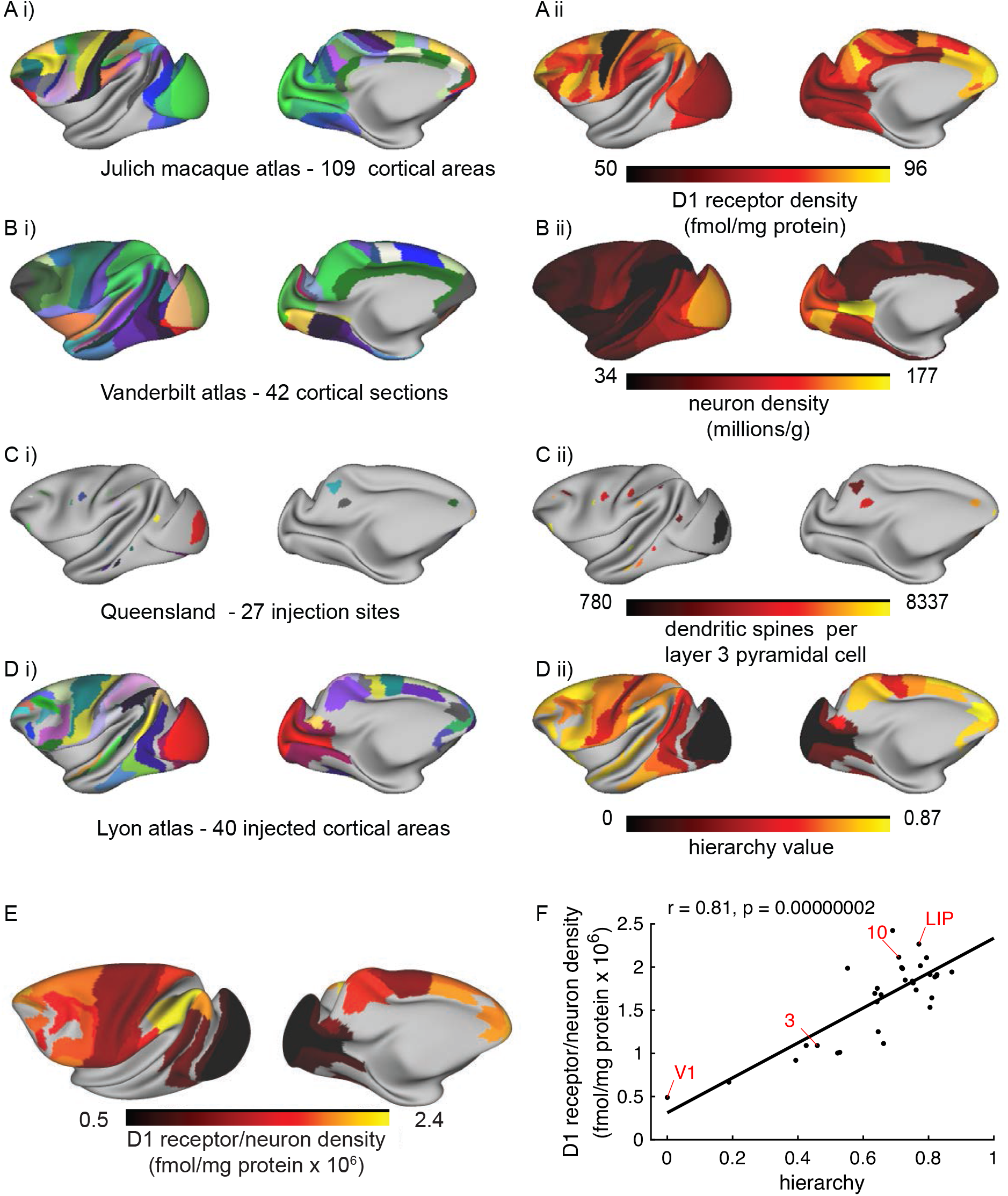
A gradient of dopamine D1 receptors per neuron across monkey cortex. A i) 109 cortical regions were identified on the basis of receptor and cytoarchitecture to create the Jülich macaque atlas. Shown here mapped onto the Yerkes19 cortical surface. A ii) D1-receptor density is low in motor and sensory cortex, and relatively high in frontal and parietal cortex. Note that the receptor density shown here does not take into account differences in neuron density across areas. B i) Collins et al., (2010) divided the entire macaque cortex into 42 slabs of tissue, which we mapped onto the Yerkes19 surface. B ii) Neuron density across cortex. C i) Injection sites for the studies of dendritic spine density by Elston and colleagues, mapped onto the Yerkes19 surface. See main text for references. C ii) Number of dendritic spines on the basal dendrites of layer III pyramidal cells. D i) 40 injected areas in the retrograde tract-tracing database of Kennedy and colleagues (Markov et al. 2014b). D ii) Cortical hierarchy, calculated based on the laminar pattern of inter-areal connections. E) The density of D1 receptors divided by neuron density. Regions that have not yet been measured shown in gray. F) There was a strong positive correlation between the D1 receptor density per neuron and the cortical hierarchy, estimated independently based on the laminar origin of cortico-cortical connections. D1R, D1 receptor density.

In order to compare the gradient of D1 receptors to other known gradients of anatomical organization in monkey cortex (Wang 2020), we carefully mapped the receptor data (Fig 1A), as well as data on neuronal density (Fig 1B) (Collins et al. 2010) and spine count (Fig 1C) (Elston 2000; Elston et al. 2001, 2005, 2011a, 2009, 2011b, 2010; Elston and Rockland 2002; Elston and Rosa 1997, 1998a,b; Elston et al. 1999) onto the Yerkes19 common cortical template, to which anatomical tract tracing data (Fig 1D i) has previously been mapped (Donahue et al. 2016).

In order to investigate a possible relationship between the pattern of D1 receptors and the cortical hierarchy, we estimated the latter using laminar connectivity data (Markov et al. 2014a). Feedforward connections tend to originate in the superficial cortical layers. In contrast, feedback connections usually originate in the deep layers (Barone et al. 2000; Felleman and Van Essen 1991; Markov et al. 2014a). If two areas are at a similar level of the hierarchy, then the connections usually arise evenly from the superficial and deep layers (Barone et al. 2000; Markov et al. 2014a). This pattern allowed us to estimate the hierarchy of 40 areas in macaque cortex (Barone et al. 2000; Markov et al. 2014a) (Fig 1D ii). This expands previous estimates of the hierarchy based on 29 or 30 areas from the same database (Markov et al. 2014a; Mejias et al. 2016).

We divided the D1 receptor density by the neuron density (Collins et al. 2010) in order to give an estimate of the D1 receptor density per neuron. D1 receptor density per neuron peaked in areas of frontal and parietal cortex, and was relatively low in early sensory cortex (Fig 1E). There was a strong positive correlation between the D1 receptor density per neuron and the cortical hierarchy (Fig 1F; *r* = 0.81*, p* = 2 × 10^−8^).

### A local cortical circuit with three types of inhibitory neurons

We built a model of a local cortical circuit which contains pyramidal cells and three types of inhibitory neurons (Fig 2A). The cortical circuit is based on a disinhibitory motif (Wang and Yang 2018; Yang et al. 2016), which was originally predicted theoretically (Wang et al. 2004a), and has since received much experimental support (Adesnik et al. 2012; Jiang et al. 2015; Pfeffer et al. 2013; Tremblay et al. 2016). We have updated details of the connectivity structure to reflect recent experimental findings (Adesnik et al. 2012; Jiang et al. 2015; Pfeffer et al. 2013; Tremblay et al. 2016). We grouped the pyramidal neurons into two separate populations. Each of these populations is selective to a particular visual feature (such as a region of visual space). Pyramidal cells excite all cell types in the circuit, with different strengths. We model two compartments in the pyramidal cells. One compartment represents the soma and proximal dendrites, and the other the distal dendrites. The dendrite is modelled as a simplified nonlinear function, adapted from Yang et al. 2016. Pyramidal cells target the soma and proximal dendrites of other pyramidal cells in the same cortical area (Kalisman et al. 2005; Markram et al. 1997; Petreanu et al. 2009). Each type of inhibitory neuron has a unique pattern of connectivity. The first inhibitory cell type targets the perisomatic area of the pyramidal cells. These cells express parvalbumin (PV) and are fast spiking (Jiang et al. 2015; Kawaguchi 1993, 1995). They are basket cells with axons that branch across wide distances, which allows them to inhibit pyramidal cells in neighboring populations (Helmstaedter et al. 2009; Kawaguchi 1995). They also inhibit other PV neurons (Jiang et al. 2015; Pfeffer et al. 2013). Compared to other inhibitory neurons, PV neurons receive a smaller proportion of excitatory inputs via NMDA receptors (Lu et al. 2007; Wang and Gao 2009). The second type of inhibitory neuron targets the distal dendrites of excitatory cells. In non-human primates, dendrite-targeting cells express calbindin (DeFelipe et al. 1989). The best characterised dendrite-targeting cell type in rodents is the Martinotti cell, which expresses somatostatin (SST/CB) (Wang et al. 2004b). These cells target all other cell types, while avoiding other Martinotti cells (Jiang et al. 2015). They also receive a strong lateral projection from pyramidal cells in neighboring columns (Adesnik et al. 2012) and receive most of their excitation via NMDA receptors (Lu et al. 2007). The third type of interneuron expresses vasoactive intestinal peptide and calretinin (VIP/CR) (Tremblay et al. 2016) and targets SST/CB inhibitory neurons (Lee et al. 2013).

**Figure 2:**
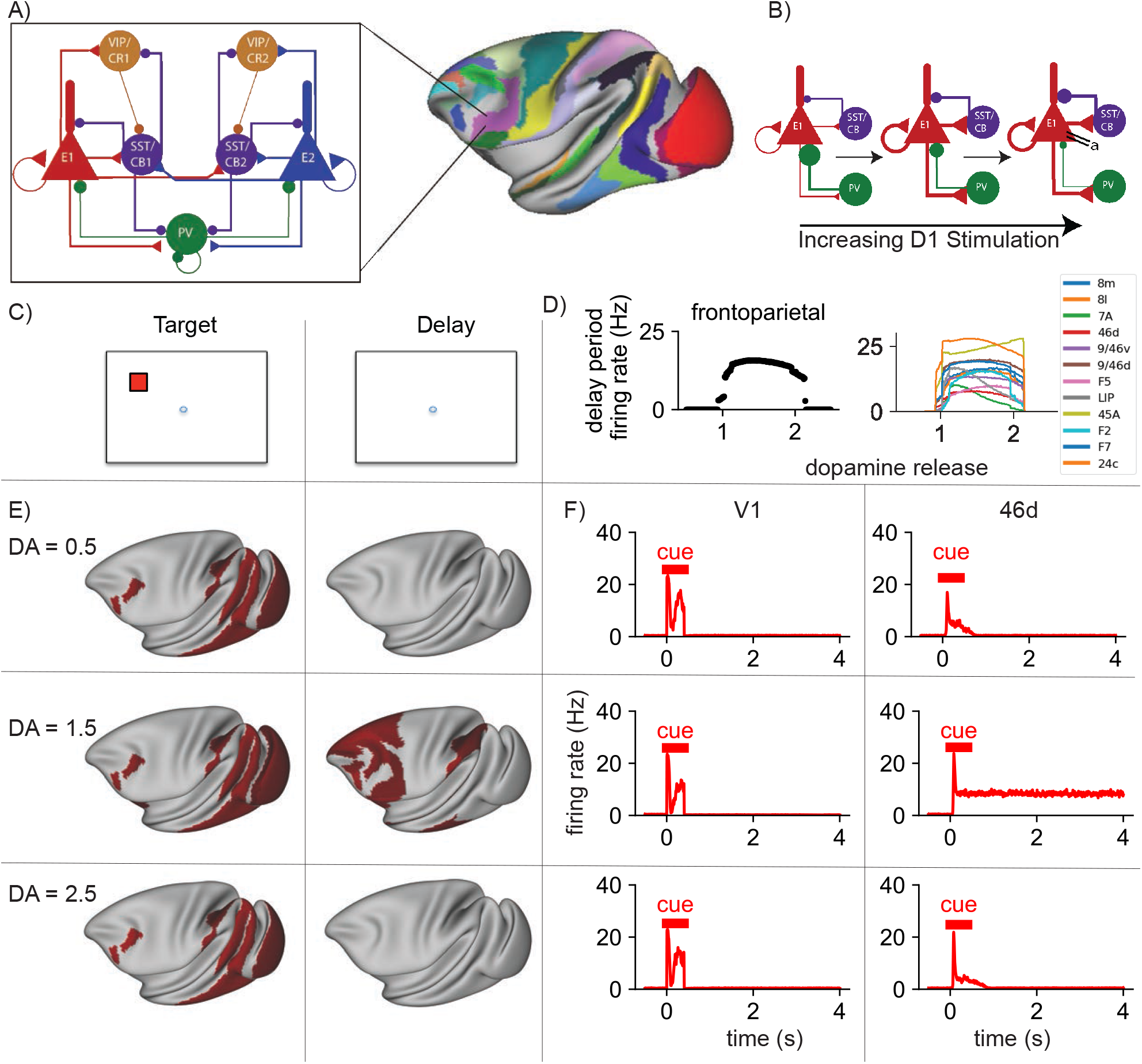
An inverted-U relationship between D1 receptor stimulation and distributed frontoparietal delay-period activity. A, left) Local circuit design. The circuit contains two populations of excitatory cells (red and blue), with each selective to a particular spatial location. The cell bodies (triangles) and dendrites (rectangles) are modeled as separate compartments. PV cells (green) inhibit the cell bodies of both excitatory populations. SST/CB cells (purple) inhibit the dendrites of the local excitatory population. VIP/CR cells (light brown) inhibit SST/CB cells. A, right) In order to construct the large-scale model, the local circuit in A) is placed at each of 40 cortical locations (shown in colours). Cortical areas differ in three properties 1) the long-range connections, 2) the spine count and 3) the dopamine D1 receptor density. B) Stimulation of D1 receptors affects the cortical circuit in three ways 1) an increase of inhibition targeting the dendrites, with a corresponding decrease in inhibition to the soma of pyramidal cells, 2) an increase in NMDA-dependent excitatory transmission for low-to-medium levels of stimulation and 3) increasing adaptation for high levels of stimulation. C) Structure of the task. The cortical network was presented with a stimulus, which it had to maintain through a delay period. D, left) Mean firing rate in the frontoparietal network at the end of the delay period, for different levels of dopamine release. There is an inverted-U relationship between dopamine release and delay period activity across the frontoparietal network. D, right) Mean delay-period activity of cortical areas as a function of dopamine release. All areas shown display persistent activity in experiments (Leavitt et al. 2017). E) Activity is shown across the cortex at different stages in the working memory task (left to right), with increasing levels of dopamine release (from top to bottom). Red represents activity in the excitatory population sensitive to the location of the target stimulus. Very low or very high levels of dopamine release resulted in reduced propagation of stimulus-related activity to frontal areas and a failure to engage persistent activity. Mid-level dopamine release enables distributed persistent activity. F) Timecourses of activity in selected cortical areas. The horizontal bars indicate the timing of cue (red) input to area V1. Activity in early visual areas such as V1 peaks in response to the stimulus, but quickly decays away after stimulus removal for all levels of dopamine release. In contrast, there is dopamine-dependent persistent activity in area 46d of prefrontal cortex. DA, cortical dopamine availability.

### Dopamine modulates interactions between multiple cell types

In our model, dopamine acted by increasing the synaptic strength of inhibition to the dendrite, and reducing the synaptic strength of inhibition to the cell body (Fig 2B) (Gao et al. 2003). In addition, dopamine increased the strength of transmission via NMDA receptors (Seamans et al. 2001a). On the other hand, high stimulation of D1 receptors resulted increased adaptation in excitatory cells (potentially an M-current, via KCNQ potassium channels, Arnsten et al. 2019), mimicking the net inhibitory effect of high concentrations of D1-agonists.

### A large-scale model of macaque cortex incorporating multiple macroscopic gradients

We then built a large-scale model of macaque cortex where the local circuit (Fig 2A, left) acted as a building block. We placed the local circuit in each of 40 cortical areas across macaque cortex (Fig 2A, right). However, these local circuits varied across areas in three key properties, namely long-distance connectivity, strength of excitation and modulation by D1 receptors. We defined the connections between areas using quantitative retrograde tract-tracing data (Markov et al. 2014b). This data was collected in the same lab under the same conditions for injections into 40 distinct cortical areas. This offers high fidelity reconstruction of the weighted and directed connections between neurons in large sections of macaque cortex (Kennedy et al. 2013). In the model, long-range connections are excitatory and target the dendrites of pyramidal cells (Petreanu et al. 2009). Long-range excitatory connections also target VIP/CR cells to a greater degree than PV or SST/CB cells (Lee et al. 2013; Wall et al. 2016). The frontal eye fields (FEF - areas 8m and 8l in the Jülich and Lyon macaque atlases) have an unusually high density of calretinin (here VIP/CR) cells (Pouget et al. 2009). To account for this, we increased the proportion of long-range input to VIP/CR cells in FEF and reduced the strength of input to the PV and SST/CB cells. We also increased the relative strength of local VIP/CR connections and reduced the relative strength of local PV connections in FEF, but found that this had no effect on model behaviour, so the simulations here are presented without the local changes in FEF.

The number of spines on the basal dendrites of layer III pyramidal neurons increases along the hierarchy (Chaudhuri et al. 2015; Elston 2007). Approximately 90% of excitatory synapses on neocortical pyramidal cells are on dendritic spines (Nimchinsky et al. 2002). On this basis, we used the spine count to modulate the strength of excitatory connections across the cortex. The strength of dopamine modulation, as described in the previous section, depended both the global dopamine release and the local D1 receptor density across cortical areas.

### An inverted-U relationship between cortical D1 receptor stimulation and distributed working memory activity

We simulated the large-scale cortical model during performance of the working memory task (Fig 2C) with different levels of dopamine release. Dopamine neurons fire bursts in response to stimuli predicting reward in working memory tasks (Schultz et al. 1993). Although striatal dopamine levels return to baseline relatively quickly following dopamine release, in the cortex dopamine levels remain elevated for seconds (Cass and Gerhardt 1995; Garris and Wightman 1994; Muller et al. 2014; Mundorf et al. 2001), which is approximately the period of one trial in our simulations. Therefore, for the majority of simulations we approximated this by setting dopamine to a constant value for each trial.

In simulations, stimulus-selective activity propagated from visual cortex to temporal, parietal and frontal cortex. Activity in visual cortex was relatively insensitive to dopamine (Fig 2E,F). In all cases, there was a strong transient response in visual areas, before a quick return to baseline firing rates, like V1 neurons recorded from behaving monkeys (Vugt et al. 2018). We observed a similar transient response in somatosensory areas in response to stimulus input to somatosensory cortex (Fig S2), as seen experimentally (Romo and Rossi-Pool 2020). Delay-period activity in a large network of prefrontal, lateral parietal and temporal areas showed an inverted-U relationship with dopamine levels (Fig 2D). A similar pattern of delay period activity was observed following somatosensory input (Fig S2). A comparable inverted-U relationship has been observed in prefrontal cortex following local application of D1 receptor agonists during working memory tasks (Vijayraghavan et al. 2007; Wang et al. 2019). A mid-range level of dopamine release engaged a distributed pattern of persistent activity throughout these areas (Fig 2E, F), but to low or too high release led to only a transient response (Fig 2F). Thus our model suggests that the inverted-U relationship between D1 receptor stimulation and persistent activity affects activity throughout a distributed fronto-parietal network, with little effect on early sensory areas.

### Long-range connectivity determines the distributed working memory activity pattern

The pattern of areas showing working memory activity in the model closely matches the areas in which persistent activity has been seen experimentally during working memory tasks. Leavitt and colleagues collated data from over 90 studies recording single-unit and multi-unit activity as monkeys performed delay tasks (Leavitt et al. 2017). For each studied area, they quantified the number of studies in which persistent activity was observed or not observed during the delay period of the task. They mapped this information onto the Lyon atlas (Kennedy et al. 2013; Markov et al. 2013, 2014a,b), which allowed us to compare our simulation with the collated delay-period activity observed in over 90 electrophysiology studies. We first divided the cortex into persistent activity and non-persistent activity areas for both the experimental data and simulation. Areas were classified in the persistent activity group if at least 3 more studies observed persistent delay period activity than a lack of such activity. We excluded areas that have been assessed in less than three studies. Areas in the simulation were classified as having persistent activity if, for the last 500ms of the trial, they had mean firing rates of at least 5Hz greater than the pre-stimulus baseline firing rates. Of the 19 cortical areas in which such activity has been assessed during the delay period in at least three papers, 18 were in agreement between the simulation and experimental results (*χ*^2^ = 15.03, *p* = 0.0001, Fig 3A). Overall, the experimentally observed persistent activity from numerous studies is reproduced, validating the model. This allows us to inspect the anatomical properties that underlie the distributed activity pattern and gain insight into the brain mechanisms that may produce it.

**Figure 3:**
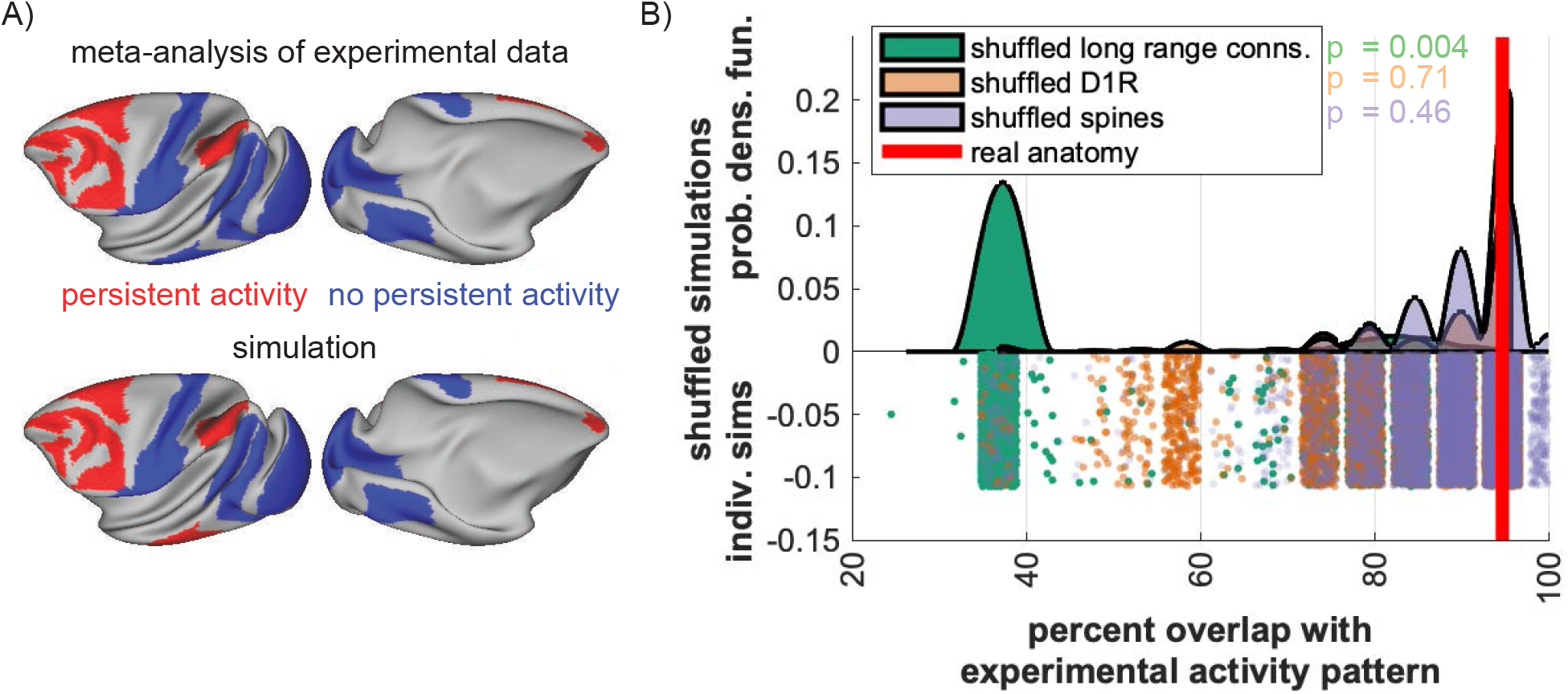
Long-range connectivity underlies the pattern of distributed working memory activity. A) There is a strong overlap (18/19 - 95%) between the pattern of persistent activity seen experimentally (Leavitt et al. 2017) and that predicted by the model. B) The percent overlap with the experimental persistent activity pattern is shown for the simulation based on the real anatomical data (red line) and for 10,000 simulations each based on shuffled long-range connections (green), shuffled D1 receptor pattern (orange) and shuffled dendritic spine counts pattern (purple). The top row half of the image shows the probability density function, and the bottom half the results of individual simulations based on shuffled anatomical data. The pattern of long-range connections was the most important determinant of the working memory activity pattern.

We repeated model simulations after shuffling the anatomical data. The delay period activity patterns for 30,000 simulations based on the shuffled anatomy were compared to the pattern observed experimentally. Ten thousand simulations were run using shuffled long-range connections, shuffled D1 receptor expression and shuffled dendritic spine expression, separately. The overlap between the experimental persistent activity pattern and the model persistent activity pattern was strongly dependent on the pattern of long-range connections (p=0.0004), but not on the pattern of D1 receptors (p = 0.71) or dendritic spine count (p = 0.46) (Fig 3B).

### Working memory deficits are more severe following lesions to areas with high D1 receptor density

Next, we examined whether lesions to individual cortical areas would affect working memory activity. The effect of lesions depended on both the area lesioned, and the level of cortical dopamine. Lesions to some cortical areas (such as areas 32, or TEO) had little effect on persistent activity in the frontoparietal network. The effect of other lesions was relatively heterogeneous. Lesions to areas LIP and 46d led to a drop in persistent activity in the remaining frontoparietal network for all dopamine levels (Fig 4A). In contrast, a lesion to area 8B led to a profound loss of frontoparietal activity for most dopamine levels, but for a restricted range, activity throughout the remaining frontoparietal network could be returned to close to the unlesioned state (Fig 4A). This suggests that for some lesions, D1 agonists or antagonists could be effective at restoring normal working memory functioning, but the correct treatment may depend on the baseline cortical dopamine levels of the patient. In contrast, other lesions seem relatively unresponsive to alterations in cortical dopamine levels.

**Figure 4:**
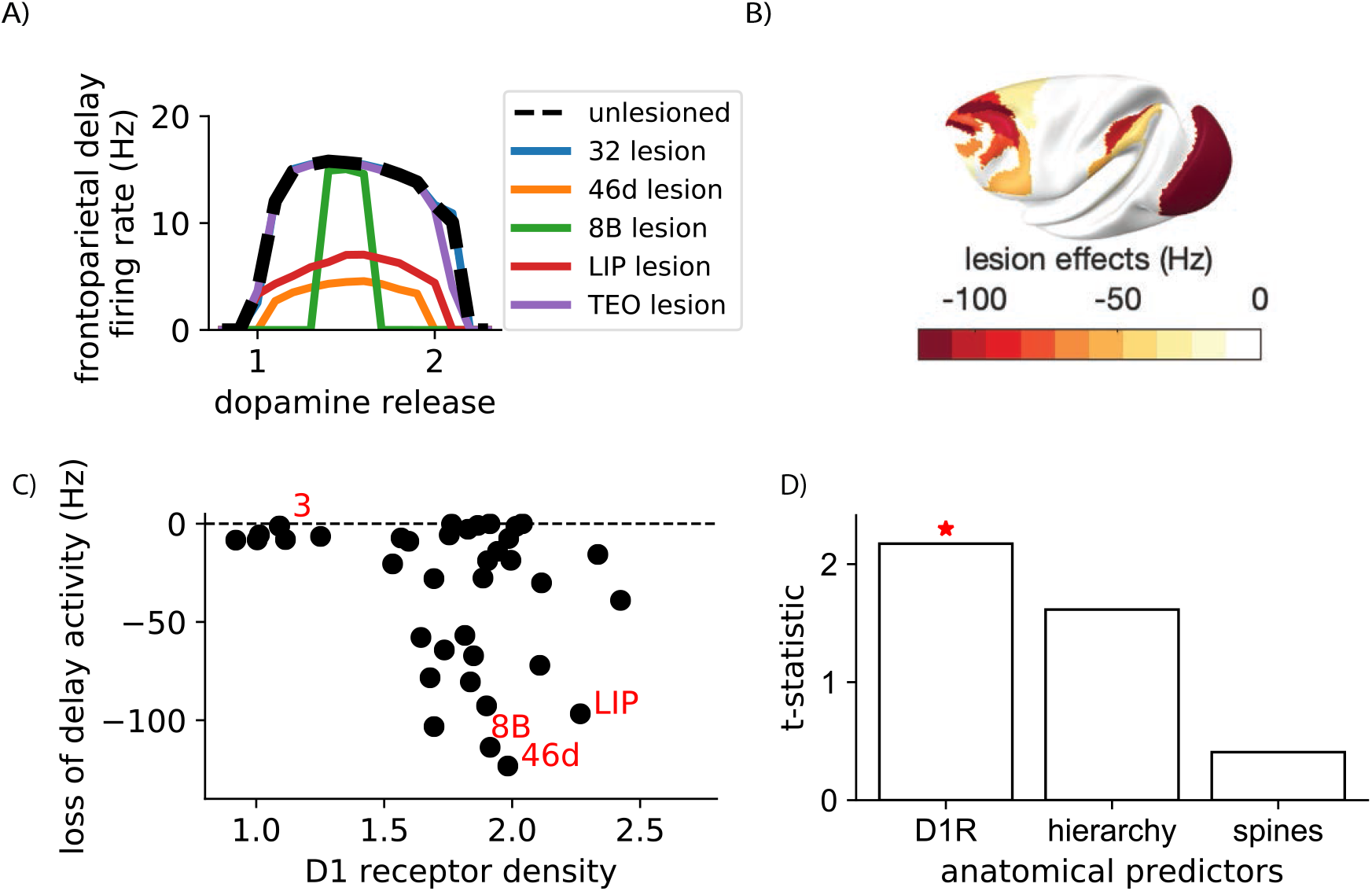
Lesions to areas with a high dopamine D1 receptor density disrupt working memory activity. A) Lesions to areas such as 46d and LIP led to reduced delay period firing across for all levels of dopamine release. Following some lesions (such as to area 8B) an optimal level of D1 receptor stimulation could restore close-to-normal working memory activity in the remaining network. B) The level of disruption to distributed working memory activity following lesions to each area, quantified as the total loss of working memory activity in the frontoparietal network summed across all dopamine release levels. Note that disruption to working memory following lesions to V1 and V2 is due to the visual stimuli being applied to V1. C) Lesions to areas with a higher D1 receptor density tended to have a larger impact on working memory activity. d) t-statistic for linear regressions predicting the drop in delay-period activity. The t-statistic for each single predictor linear regression model is shown separately. DA, dopamine release. D1R, D1 receptor density.

Lesions to area V1 and V2 led to a complete loss of visual working memory activity (Fig 4B). However, this was due to the fact that a visual stimulus must go through area V1 in order to gain access to the working memory system. We confirmed this by showing that lesions to V1 and V2 had no effect on working memory when somatosensory stimuli were used (with stimulus presented to primary somatosensory area 3). In the somatosensory working memory task, lesions to early somatosensory areas, and frontoparietal network areas caused memory deficits (Fig S3). This clearly separates early sensory areas, which are required for signal propagation to the working memory system, from core cross-modal working memory areas in prefrontal and posterior parietal cortex.

We quantified the effect of lesions as the difference between the inverted-U curved for the unlesioned and lesioned networks (Fig 4A). In this way, the lesions to area 8B and 46d have similarly large effects on average across all dopamine levels (Fig 4B,C). We then tested whether the anatomical data (namely the D1 receptor density, the spine count or the cortical hierarchy) could predict the effects of cortical lesions on working memory activity. D1 receptor density (F = 4.72, p = 0.036, Fig 4C) was the strongest predictor of the lesion effects, and adding hierarchy or spine count to the model did not significantly improve the fit (Fig 4D). Thus, lesions to areas with a higher D1 receptor density are more likely to disrupt working memory activity.

### Dopamine shifts between activity-silent and persistent activity modes of working memory

We endowed the large-scale model with short-term plasticity in order to assess the possibility of activity-silent working memory in the large-scale network. Short-term plasticity was implemented at all synapses between excitatory cells (Hempel et al. 2000; Wang et al. 2004b) (using the same parameters as Mongillo et al. 2008), and from excitatory to SST/CB cells (Lee et al. 2013; Silberberg and Markram 2007). We investigated activity-silent representations by ‘pinging’ the system, with a neutral stimulus, and reading out the activity generated in response, similar to the experimental protocol in Wolff et al. 2017 (Fig 5A i).

**Figure 5:**
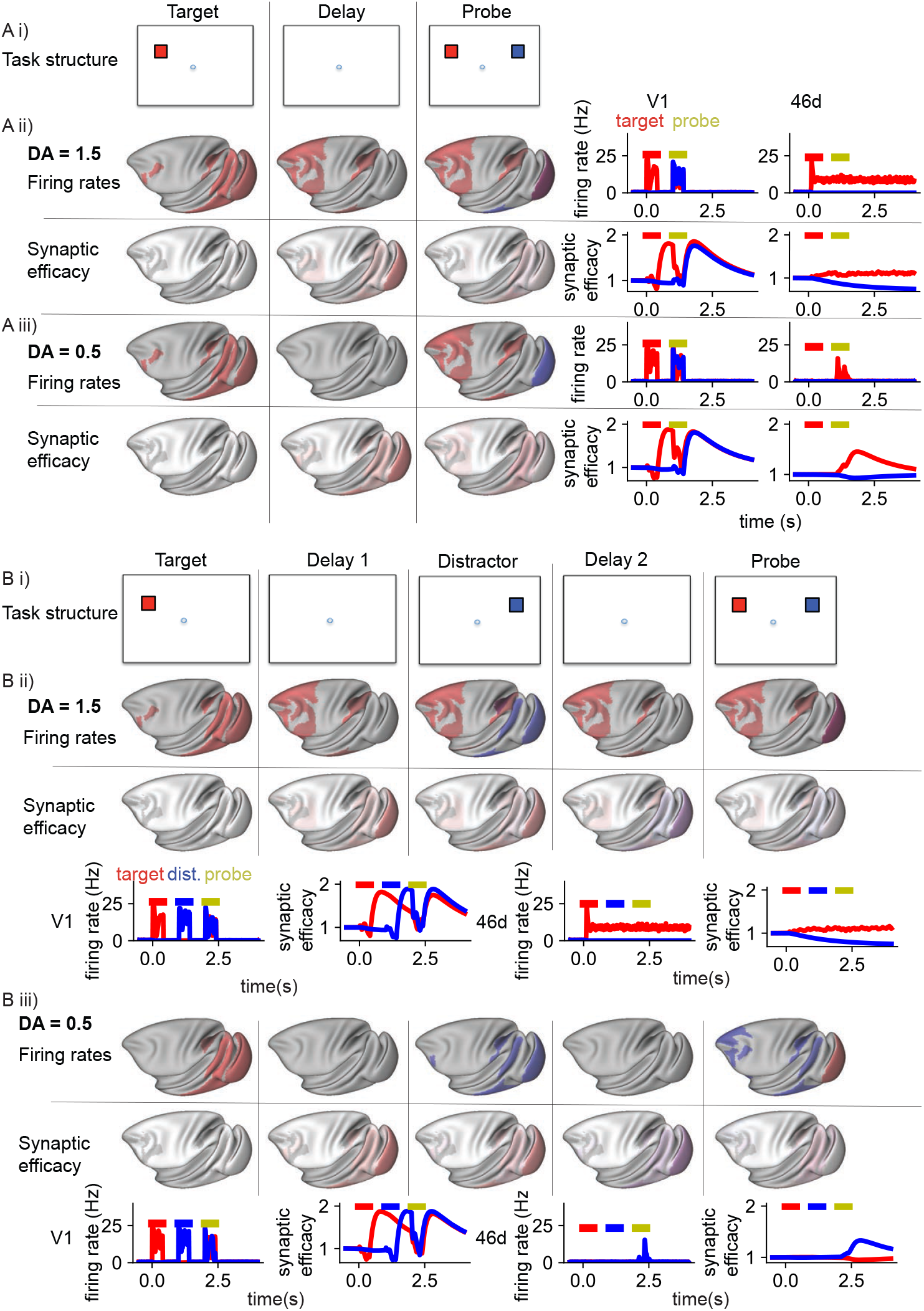
A dopamine-dependent shift between distractible activity-silent and distractor-resistant persistent-activity states. A i) Task structure. A target stimulus was followed by a delay and a probe stimulus. A ii) For mid-level dopamine release, activity relating to the target stimulus propagated from V1 through the hierarchy, and was maintained in persistent activity throughout the frontoparietal network. Top: firing rates on the surface (left) and in selected areas (right). Bottom: synaptic efficacy. A iii) For low-level dopamine release, activity (Top) in response to the stimulus was transient in visual and some frontoparietal areas. There was no persistent activity through the delay period. However, in response to the probe stimulus, activity representing the original target stimulus was regenerated throughout frontoparietal cortex. Bottom: The memory for the stimulus was stored as a short-term plasticity-dependent increase in synaptic efficacy through the delay period. This was particularly prominent for synapses from neurons with their cell bodies in sensory areas. B i) Task structure. A target stimulus was followed by a delay period, a distractor, another delay period and a probe stimulus. B ii) For mid-level dopamine release, target-related activity was maintained in persistent activity throughout the frontoparietal network, throughout the delay period through the distractor until the end of the trial. B iii) For low-level dopamine release, frontoparietal activity related to the most recent stimulus (i.e. the distractor) was regenerated during this probe stimulus. DA, dopamine release.

For optimal mid levels of dopamine release (Fig 5A ii), the model generated persistent activity that was very similar to the network without short-term plasticity. For both low and high levels of dopamine release there was no persistent activity (Fig 5 A iii). However, when we ‘pinged’ the system with a neutral stimulus, activity relating to the target cue was transiently generated throughout the frontoparietal network Fig 5 A iii). Short-term synaptic plasticity increased the synaptic efficacy for connections between neurons coding for the target stimulus during the delay period. However, most of the increase in synaptic efficacy was in synaptic connections from neurons in sensory areas (Fig 5 A iii). When short-term synaptic plasticity was restricted to neurons only in sensory areas, pinging the system still resulted in a reactivation of the target-related activity (Fig S4). This suggests that synaptic plasticity in local prefrontal cortical neurons is not required for activity-silent working memory. However, when we restricted short-term plasticity to only local connections, we could not regenerate activity related to the target stimulus with the ping. This suggests that short-term plasticity on long-range connections from sensory areas is required for activity-silent working memory in the large-scale cortical network. Furthermore, dopamine release can switch the system from an activity-silent, to a persistent activity regime.

Why does the brain have two parallel systems for holding items in short-term memory? To explore this further, we simulated the model using a “ping protocol” (Rose et al. 2016; Wolff et al. 2017). After a behaviorally relevant cue (stimulus A) and during the delay period, we introduced a distractor (stimulus B) which should be filtered out by the network; then a neutral ping stimulus exciting both neural populations is presented (Fig 5B i). For mid-level dopamine release, persistent activity coding for the target stimulus is engaged, and maintained through the distractor and ping (Fig 5B ii). The distractor is only transiently represented in IT and LIP (compare with Suzuki and Gottlieb 2013), but does not reach most of the frontoparietal network. In the low and high dopamine cases, during the ping, activity-silent mechanisms regenerate activity related to the last encoded stimulus, namely the distractor, in frontal and parietal cortex (Fig 5B iii). This suggests that dopamine can switch the cortex from an activity-silent working memory mode to a robust, distractor-resistant persistent activity working memory mode.

### Dopamine increases distractor resistance by shifting the subcellular target of inhibition

How does dopamine protect working memory from distraction? To examine this question, we analysed activity within VIP/CR and SST/CB neurons during a working memory task with a distractor (Fig 6A). SST/CB and VIP/CR neurons are in competition, as they mutually inhibit each other. When SST/CB cell firing is higher, the pyramidal cell dendrites are relatively inhibited; conversely when VIP/CR cell firing is higher, the pyramidal cell dendrites are disinhibited. Each cortical area in the model contains two selective populations of pyramidal, SST/CB and VIP/CR cells. We first analysed trials in which the model successfully ignores the distractor. In the target-selective populations, the VIP/CR neurons fire at a much higher rate than the SST/CB neurons (Fig 6B, C). Thus the dendrites of the pyramidal cells sensitive to the target are disinhibited, allowing long-range target-related activity to flow between cortical areas. In the distractor senstitive populations, throughout the frontoparietal network, the SST/CB neurons fire at a slightly higher rate than the VIP/CR cells. Thus distractor-related activity from other cortical areas is blocked from entering the dendrites of distractor-sensitive pyramidal cells in frontal and parietal cortex.

**Figure 6:**
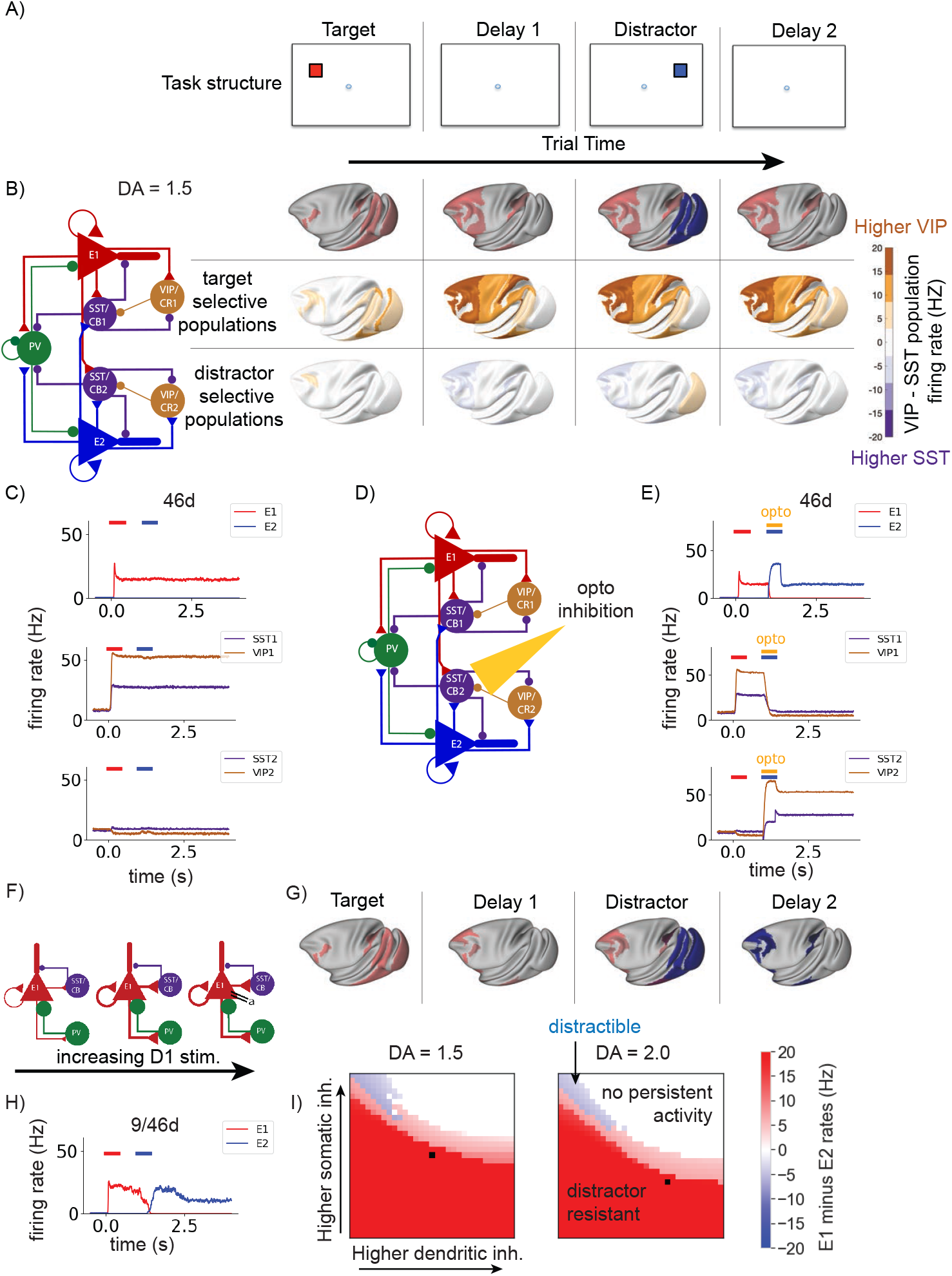
Dopamine increases distractor resistance by shifting the subcellular target of inhibition. A) Task structure. A target stimulus was followed by a delay, a distractor stimulus and another delay period. B) For mid-level dopamine release, persistent target-related activity (red) was present in the frontoparietal network through the delay and the distractor until the end of the trial. Each cortical area contains populations of excitatory, somatostatin and VIP/CR cells that respond to the target stimulus (E1, SST1, VIP1), separate populations sensitive to the distractor stimulus (E2, SST2, VIP2) and PV cells. B and C) Throughout the delay period and distractor stimulus, activity in VIP1 is higher than in SST1, leading to disinhibition of the E1 dendrite. In contrast, activity in VIP2 is slightly lower than in SST2, leading to inhibition of the E2 dendrite. D) We tested the causal effect of the SST2 activity, by transiently inactivating SST2 populations in the frontoparietal network during the presentation of the distractor stimulus (simulating optogenetic inhibition). E) On trials in which SST2 populations were inhibited, activity relating to the distractor stimulus propagated from early sensory areas to frontoparietal areas, and the network maintainted distractor-related activity until the end of the trial. F) We then removed the dopamine modulation of somatic and dendritic inhibition, while keeping the other dopamine effects as before. G,H) Without the dopamine-dependent switch towards dendritic inhibition, the network became distractible, with distractor-related activity dominating at the end of the trial. I) We identified the model behaviour for different dopamine levels, across different levels of dendritic and somatic inhibition. Consistently across dopamine levels, higher somatic, and lower dendritic inhibition was associated with distractible working memory (blue). In contrast, lower somatic, and higher somatic inhibition was associated with distractor-resistant working memory (red). High dendritic and high somatic inhibition results in no persistant activity (white). The levels of dendritic and somatic inhibition associated with the standard dopamine modulation used in the rest of the paper marked by a black square. DA, cortical dopamine availability.

The relative inhibition of distractor-population dendrites is rather subtle. To test the importance of this effect, we simulated optogenetic inhibition of the SST/CB populations that inhibit the distractor-sensitive pyramidal cells (SST/CB2, Fig 6D). We transiently inhibited the SST/CB2 cells in the frontoparietal network during the presentation of the distractor. This transient inhibition of SST/CB2 cells was sufficient to switch the network to a distractible state, with the distractor stimulus held in working memory until the end of the trial (Fig 6E).

As dopamine increases the strength of inhibition to the dendrites, and decreases inhibition to the soma, it is possible that this aspect of dopamine modulation enhances distractor-resistance of the system. We removed this effect of dopamine modulation, while applying the other effects of dopamine as before (Fig 6F). We repeated the working memory task with the distractor with a mid-level of dopamine, which normally results in distractor-resistant working memory (Fig 6 A,B). Without the shift of inhibition from the soma to the dendrite, the system becomes distractible (Fig 6G, H). We searched the parameter space, and found that, when local cortical areas were independently capable of maintaining persistent activity (e.g., *μ*^*E,E*^ = 1.25, 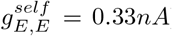), high somatic inhibition and low dendritic inhibition was generally associated with distractibility (Fig 6I). Low somatic and high dendritic inhibition was associated with distractor-resistant behaviour (Fig 6I, S5). Therefore dopamine shifts inhibition from the soma to the dendrite, and stops distractor stimuli from sensory areas disrupting ongoing persistent activity in the frontoparietal network.

### Learning to optimally time dopamine release via reinforcement

We have shown how dopamine can render the cortex resistant to distraction, but one potential objection is that a behavioral relevant cue does not always precede a distractor. As a matter of fact, in real life we experience a constant flow of sensory inputs, our working memory system must be flexible in determining the timing of relevant versus irrelevant information. In laboratory experiments, one can assess this flexibility by presenting stimuli to be ignored before the relevant stimulus appears (e.g. Atlan et al. 2018). Dopamine neurons fire in response to task-relevant stimuli (Schultz et al. 1993), but should not fire in response to task-irrelevant distracting stimuli. We hypothesised that the correct timing of dopamine release could be learned by simple reward-learning mechanisms.

We added a simplified model of the ventral tegmental area (VTA) with GABAergic and dopaminergic neurons to our large-scale cortical model (Fig 7A). Cortical pyramidal cells target both GABAergic and dopaminergic cells in the VTA (Soden et al. 2020). Dopaminergic cells are also strongly inhibited by local VTA GABAergic cells (Soden et al. 2020). Dopamine is released in cortex in response to VTA dopaminergic neuron firing, and cortical dopamine levels slowly return to baseline following cessation of dopaminergic neuron firing (Cass and Gerhardt 1995; Garris and Wightman 1994; Muller et al. 2014; Mundorf et al. 2001). The synaptic strengths of cortical inputs from the selected populations to VTA populations are increased following a reward, and weakened following an incorrect response (Harnett et al. 2009; Soltani and Wang 2006).

**Figure 7:**
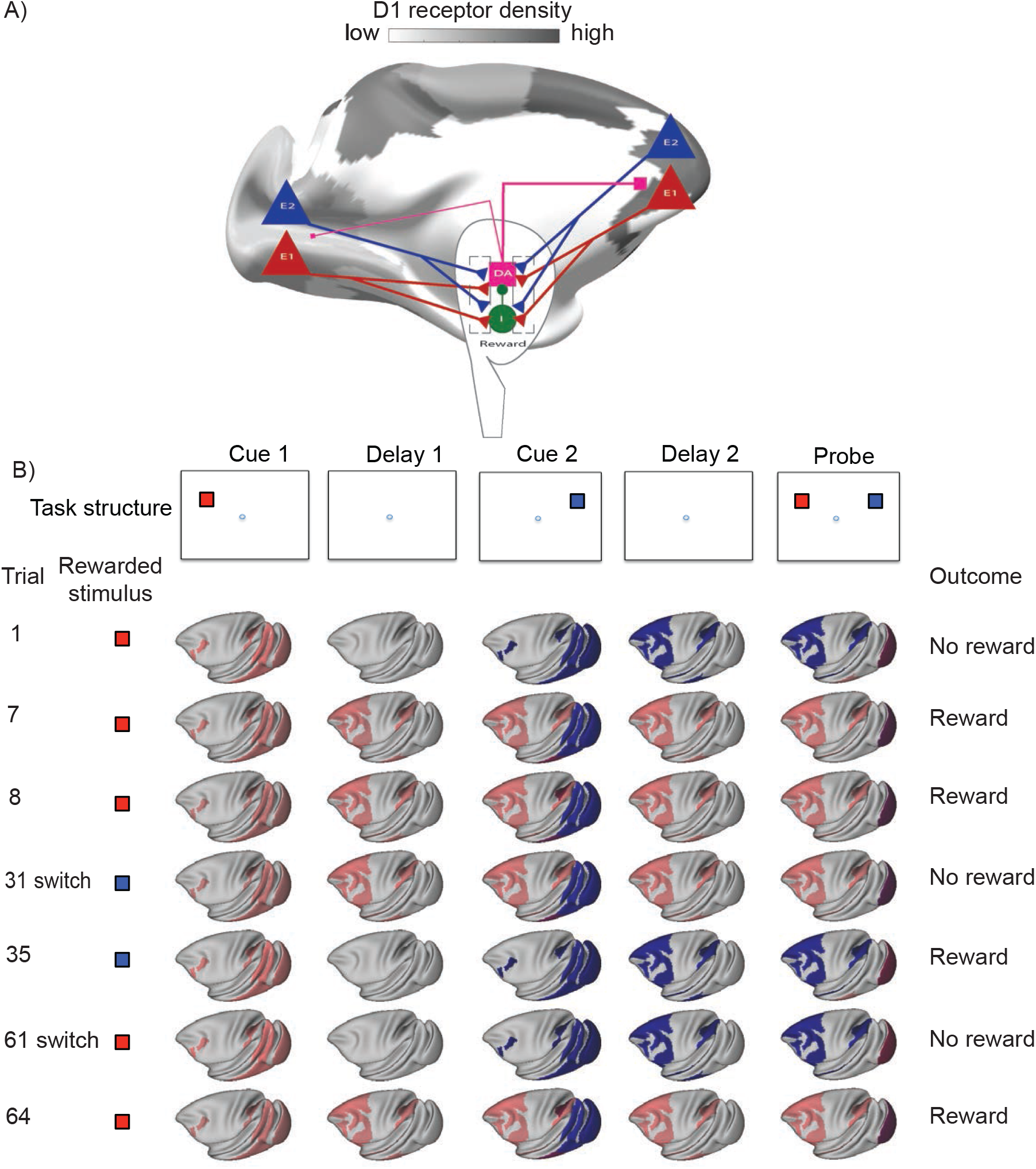
Reward-dependent learning of dopamine release appropriately engages persistent activity mechanisms. A) We designed a simplified VTA model and connected this bidirectionally to the large-scale cortical model. The VTA contained dopaminergic and GABAergic neuron populations. Dopamine was released dynamically depending on dopaminergic neuron activity. The strength of cortical inputs to VTA dopaminergic and GABAergic cells was updated at the end of each trial on the basis of trial outcome and choice. B) We simulated a task with two cues (red and blue) followed by a probe stimulus. The rewarded stimulus changed every 30 trials. Following each switch, after a few trials the network learns to store the appropriate stimulus in distributed persistent activity. This depends on high dopamine release in response to the rewarded stimulus and low release in response to the unrewarded stimulus.

We tested the model on the target-distractor-ping task introduced earlier (Fig 5B i; 7 B). For the first 30 trials, the first stimulus (Cue 1, red) was rewarded (rule 1). For the following 30 trials the second stimulus (Cue 2, blue) was rewarded (rule 2). For the final 30 trials, we switched back to rule 1 (Fig 7B). By the seventh trial of the first block distractor-resistant persistent activity emerges, and the first cue is correctly remembered. This behaviour remained until the next block. Following a few trials of the second block, dopamine release in response to the first stimulus was reduced, and neural populations throughout the cortex only transiently represented the first (now irrelevant) stimulus. However, dopamine response to the second stimulus increased, so that persistent activity was engaged following the second stimulus. Following the second rule switch, the system again switched back to engaging persistent activity in response to the first cue. Additionally, the number of trials to a successful switch gradually decreased with each switch. We further tested the model on a version of the task in which the relevant red cue could be shown either first or second within a block, before the blue cue became relevant in the second block. The model was also able to learn this task, although it took more trials (10-15) to learn the switch (for the first few blocks). Thus, with simple reward-learning mechanisms, the optimal timing of dopamine release can be learned, allowing flexible engagement of distributed persistent activity in working memory.

## Discussion

Traditional neural network models of working memory and dopamine have focused on simulating activity within a local cortical circuit (Brunel and Wang 2001; Durstewitz et al. 2000). Here, by combining large-scale systematic receptor anatomy with large-scale cortical modelling, we show that dopamine can engage distributed persistent activity across multiple cortical areas underlying conscious, active working memory. The discovery of a macroscopic gradient of dopamine D1 receptor density per neuron, reported here for the first time, enabled us to investigate dopamine modulation across the large-scale primate cortex in a connectome- and biophysically-based model. The model represents a cross-level computational platform, endowed with multiple cell types. Our work leads to new predictions that would not have been possible with local circuit models. First, filtering out distractors to ensure robust working memory depends on dopamine action by virtue of shifting subcellular inhibition in favor of input control at dendrites of pyramidal neurons, and plasticity in the dopamine system can flexibly learn what stimuli are behaviorally relevant in a sequence of stimuli occurring in time. Second, when a short-term memory trace is encoded by an active-silent synaptic mechanisms, we found that short-term synaptic plasticity of long-range cortical connections is more important than local connections. ‘Activity silent’ memory strength is always the strongest for the latest stimulus, so this cannot subserve working memory in the presence of distractors. Third, the effect of lesioning a cortical area depends not only on that area’s connectivity with the rest of the system but also the strength of its dopamine modulation.

### A gradient of D1 receptors along the cortical hierarchy

Dopamine exerts a powerful influence on cortical computations across a range of cognitive functions (Brozoski et al. 1979; Goldman-Rakic 1995). By quantifying the D1 receptor density across cortical areas, we can identify the limits to which dopamine can modulate cortical activity. In order to create a high-resolution, and high-fidelity map of cortical dopamine receptor architecture, we used quantitative *in-vitro* receptor autoradiography (Zilles and Palomero-Gallagher 2017). PET and SPECT scans provide the advantages of *in-vivo* measurements, such as information on individual and group differences, but are limited in spatial resolution and signal-to-noise ratio (Cassidy et al. 2016; Cools and D’Esposito 2011; Froudist-Walsh et al. 2017a; Laruelle et al. 1996; Roffman et al. 2016) and are often unreliable for cortical measurements (Egerton et al. 2010; Farde et al. 1988). Gene expression methods have certain advantages, especially RNA sequencing which can provide cell-specific data. However, mRNA expression is not always closely related to, or even positively correlated with the receptor density at the synapse (Arnatkeviciute et al. 2019; Schwanhäusser et al. 2011). Receptor density at the synapse is the functionally important quantity, and is directly measured here. The map of D1 receptor density here greatly expands previous descriptions of D1 receptor densities (Goldman-Rakic et al. 1990; Impieri et al. 2019; Lidow et al. 1991; Niu et al. 2020; Richfield et al. 1989). We show that D1 receptor density increases along the cortical hierarchy, peaking in prefrontal and posterior parietal cortex. This gradient of dopamine receptors is an anatomical basis by which dopamine can modulate higher cognitive processing.

### An inverted-U relationship between dopamine and distributed working memory activity

Previous experimental and modelling studies have shown an inverted-U relationship between D1 receptor stimulation and persistent activity in the prefrontal cortex in monkeys performing working memory tasks (Brunel and Wang 2001; Vijayraghavan et al. 2007; Wang et al. 2019). Stimulation of VTA (presumably leading to cortical dopamine release) in resting monkeys also has an inverted-U effect on cortical activity in distributed areas across cortex (Murris et al. 2020). By constructing a novel-large scale model based on the D1 receptor map and tract-tracing data, we were able to show that the inverted-U relationship between D1 receptor stimulation and persistent activity held across frontal and parietal cortex during working memory. A network of frontal and parietal areas engaged in persistent activity together within a wide range of D1 receptor stimulation, with some variability in the degree to which D1 receptor stimulation affected persistent activity within areas. The working memory activity pattern was strikingly similar to that seen experimentally, according to a meta-analysis of 90 electophysiology studies of delay period activity in monkey cortex (Leavitt et al. 2017). By analysing the model, we found that the pattern of long-range connections was the strongest determinant of the pattern of working memory activity.

### Lesions to areas with a high D1 receptor density disrupt working memory

Working memory activity was disrupted most strongly by lesions to areas with a high D1 receptor density, a prediction that can be tested experimentally. Human patients with traumatic brain injury often have working memory deficits (Dunning et al. 2016). Pharmacological treatment of these deficits, including with dopaminergic drugs, has seen mixed success (Froudist-Walsh et al. 2017b). Our model simulations suggest that for lesions to some cortical areas D1 agonists or antagonists could be effective at restoring normal working memory functioning, but the correct treatment may depend on the baseline cortical dopamine levels of the patient. In the future, our model could be adapted to simulate the working memory deficits and potential treatments of neurology or psychiatry patients based on their particular anatomy and patterns of cortical dopamine release or receptor density (Abi-Dargham et al. 2002; Cassidy et al. 2016; Pettersson-Yeo et al. 2011; Slifstein et al. 2015).

### Ignition, silent activity and maintenance

The strong and distributed activation of frontal and parietal cortex is reminiscent of the ignition response to consciously observed stimuli (Dehaene and Changeux 2011; Dehaene et al. 1998, 2003; Vugt et al. 2018). Conscious ignition and working memory maintenance have similar spatial patterns of activity (Trübutschek et al. 2017), and it has been suggested that conscious ignition is a first step to the entry of information to working memory (Mashour et al. 2020). However, for very low or high levels of D1 receptor stimulation, it was possible to maintain stimulus information in the absence of persistent activity, via synaptic mechanisms. This was possible regardless of whether the original stimulus resulted in prefrontal activity. Unconscious working memory results in similarly correct behavioural performance, without the characteristic frontoparietal delay-period activity (Trübutschek et al. 2017, 2019). Unconscious working memory is thought to rely on ‘activity-silent’ synaptic mechanisms (Stokes 2015; Trübutschek et al. 2017). Slow cellular and synaptic processes may not only contribute to such mechanisms but also induce trial-by-trial history-dependent effects (Barbosa et al. 2020; Bliss and D’Esposito 2017; Carter and Wang 2007; Pereira and Wang 2015). Previous models of working memory with short-term synaptic plasticity have focused on local activity in the prefrontal cortex (Mongillo et al. 2008; Stokes 2015), and thus implicitly imply that it is short-term plasticity in local connections between prefrontal neurons that stores the memory trace. These models also assume that prefrontal neurons must have been sufficiently activated by the original stimulus in order to enable local short-term facilitation, a proposition that seems inconsistent with unconscious working memory. We show that short-term facilitation in long-range feedforward connections from early sensory areas to frontal and parietal cortex is a potential substrate for ‘activity-silent’ working memory in the absence of an initial prefrontal response to the stimulus. Given that experimental and modelling evidence suggests that manipulation of stored information requires a re-emergence of strong distributed activity (Masse et al. 2019; Trübutschek et al. 2019), the silent state may be better described as ‘short-term memory’, as noted by other authors (Mashour et al. 2020; Masse et al. 2020). The brain may then reserve widespread persistent activity for important information that must be used and manipulated to drive behavior. The model in Barbosa et al. 2020 suggests nonspecific excitatory or inhibitory currents could account for switches between active and silent states. We propose that dopamine could in fact account for the switch from silent to active state. Indeed, due to the inverted-U relationship between dopamine and persistent firing, a dopamine response to the reward at the end of a trial could also terminate persistent activity. Our model also suggests that memories stored in the active state are more robust to distraction compared to memories stored in the silent state. This suggests that dopamine may be released in order to focus attention on salient items in working memory, and protect them from distraction.

### Dopamine increases distractor resistance by shifting the subcellular target of inhibition

The resilience of the active working memory state in the model depended on SST/CB cells blocking distracting inputs from sensory areas to the dendrites of pyramidal cells in frontal and parietal cortex. Previous modelling work on local cortical circuits has suggested that greater dendritic and less somatic inhibition could increase distractor-resistance (Wang et al. 2004a), and that selective disinhibition of the dendrite (through VIP/CR cells) could selectively allow information to be passed through the network (Yang et al. 2016). In our large-scale model, VIP/CR cells selectively disinhibited the dendrites of cells selective to the target stimulus, allowing target-related activity to flow through the cortical network. D1 receptors in monkey cortex are more strongly expressed on SST/CB neurons than other interneuron types (Mueller et al. 2019), and application of dopamine to a frontal cortex slice increases inhibition to the dendrite, and decreases inhibition to the soma of pyramidal cells (Gao et al. 2003). We found that, as long as local cortical areas (or potentially cortico-subcortical loops) are capable of maintaining persistent activity, then shifting the balance of inhibition from the soma to the dendrite can allow for maintenance of an active representation of a stimulus in persistent activity, while shielding it from distracting input from sensory areas.

Distractor-resistance in response to all stimuli could render the working memory system inflexible, and unresponsive to new, potentially important inputs. Inspired by previous models of prefrontal cortex and basal ganglia (Braver and Cohen 2000; Frank 2005), we show that using simple reward-based learning, the timing of dopamine release to the cortex can be learned in order to engage distributed persistent activity throughout the frontoparietal network in response to behaviourally-relevant stimuli. In contrast, irrelevant, or less salient stimuli result in lower dopamine release, and may be remembered via silent mechanisms, or forgotten.

## Conclusion

We uncovered a macroscopic gradient of dopamine D1 receptor density along the cortical hierarchy. By building a novel large-scale anatomically-constrained model of monkey cortex, we show how dopamine can engage robust distributed persistent activity mechanisms across connected higher cortical areas, and protect memories of behaviourally relevant-stimuli from distraction. As distributed persistent activity is necessary for the manipulation of thoughts in working memory (Masse et al. 2019; Trübutschek et al. 2019), dopamine release in the cortex may be a key step towards higher cognitive thought.

## Methods

### Overview of anatomical data

In this study, we combine post-mortem anatomical data on receptor densities, white matter connectivity, neuron densities and dendritic spine counts. Each of these four anatomical measures was originally quantified using different parcellations of cortex. Large sections of the temporal lobe are not yet quantified for either the receptor autoradiography data, or the tract-tracing connectivity data. Collection of this data is underway and will be made available in future studies. With the exception of the receptor densities in the superior parietal lobe (Impieri et al. 2019) and intraparietal sulcus (Niu et al. 2020), all D1 receptor densities are reported for the first time in this study.

### A note on notation

Subscripts in square brackets, such as [*k*] are used to denote cortical areas themselves. Subscripts not in brackets, such as *i* are used to denote populations of neurons within a cortical area. Superscripts are used to provide further clarifying information. We use the convention that targets are listed before sources, so that *g*_*i,j*_ would denote the strength of a connection from neural population j to neural population i. Parameter values are listed in Table 4.

### Quantification of receptor density across cortex - in-vitro autoradiography

We analysed the brains of three adult male *Macaca fascicularis* specimens (between 6 and 8 years old; body weight between 5.2 and 6.6 kg) obtained from Covance, Münster, where they were used as control animals for pharmaceutical studies performed in compliance with legal requirements.

All experimental protocols were in accordance with the guidelines of the European laws for the care and use of animals for scientific purposes. Animals were sacrificed by means of an intravenous lethal dose of sodium pentobarbital. Brains were removed immediately from the skull, and brain stem and cerebellum were dissected off in close proximity to the cerebral peduncles. Hemispheres were separated and then cut into a rostral and a caudal block by a cut in the coronal plane of sectioning between the central and arcuate sulci. These blocks were frozen in isopentane at −40C to −50C, and then stored in airtight plastic bags at −70C. Each block was serially sectioned in the coronal plane (section thickness 20 *μm*) using a cryostat microtome (CM 3050, Leica, Germany). Sections were thaw-mounted on gelatine-coated slides, freeze-dried overnight and processed for visualization of D1 or D2 receptors, cell bodies (Merker 1983) or myelin (Gallyas 1979). Quantitative *in-vitro* receptor autoradiography was applied to label dopaminergic D1 and D2 receptors according to previously published protocols (Palomero-Gallagher and Zilles 2018; Zilles et al. 2002) encompassing a preincubation, a main incubation and a final rinsing step. For visualization of the D1 receptor, sections were first rehydrated and endogenous substances removed during a 20 minute preincubation at room temperature in a 50 mM Tris-HCl buffer (pH 7.4) containing 120 mM NaCl, 5 mM KCl, 2 mM CaCl_2_ and 1 mM MgCl_2_. During the main incubation, sections were incubated with either 0.5 nM [^3^H]SCH 23390 alone (to determine total binding), or with 0.5 nM [^3^H]SCH 23390 and 1 mM of the displacer mianserin (to determine the proportion of displaceable, non-specific binding) for 90 minutes at room temperature in the same buffer as used for the preincubation. Finally, the rinsing procedure consisted of two 20 minutes washing steps in cold buffer followed by a short dip in distilled water. For visualization of the D2 receptor, sections were preincubated 50 mM Tris-HCl buffer (pH 7.4) containing 150 mM NaCl and 1% ascorbate. In the main incubation, sections were incubated with either 0.3 nM [^3^H]raclopride alone, or with 0.3 nM [^3^H]raclopride and 1 *μ*M of the displacer 1 *μ*M butaclamol for 45 minutes at room temperature in the same buffer as used for the preincubation. Rinsing consisted of six 1 minute washing steps in cold buffer followed by a short dip in distilled water. Specific binding is the difference between total and non-specific binding. Since the ligands and binding protocols used resulted in a displaceable binding, which was less than 5% of the total binding, total binding is considered to be equivalent of specific binding. Sections were dried in a cold stream of air, exposed together with plastic scales of known radioactivity against tritium-sensitive films (Hyperfilm, Amersham) for six (for the D1 receptor) or eight (for the D2 receptor) weeks, and ensuing autoradiographs processed by densitometry with a video-based image analysing technique (Palomero-Gallagher and Zilles 2018; Zilles et al. 2002). Autoradiographs were digitized using a CCD-camera, and stored as 8-bit grey value images with a spatial resolution of 2080×1542 pixels. Grey values (*g*) in the co-exposed scales as well as experimental conditions were used to create a regression curve with which grey values in each pixel of an autoradiograph were transformed into binding site densities (Bmax) in fmol/mg protein by means of the formula

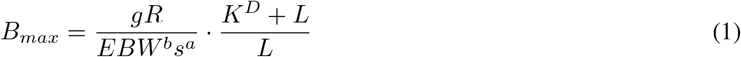

where *R* is the radioactivity concentration (cpm) in a scale, *E* the efficiency of the scintillation counter used to determine the amount of radioactivity in the incubation buffer, *B* the number of decays per unit of time and radioactivity, *W*_*b*_ the protein weight of a standard, *S*_*a*_ the specific activity of the ligand, *K*_*D*_ the dissociation constant of the ligand, and *L* the free concentration of the ligand during incubation. For visualization purposes solely, autoradiographs were subsequently pseudo-colour coded by linear contrast enhancement and assignment of equally spaced density ranges to a spectral arrangement of eleven colours.

Cortical areas were identified by cytoarchitectonic analysis and receptor densities measured at comparable sites in the adjacent sections processed for receptor visualization. The mean receptor density for each area over a series of 3–5 sections per animal and receptor was determined by density profiles extracted vertical to the cortical surface using Matlab-based in house software (Palomero-Gallagher and Zilles 2018).

### Neuronal density data

The *in-vitro* autoradiography data accurately quantifies the density of receptors across cortex. However, it is important to bear in mind that the density of neurons also varies across the cortex. Collins and colleagues measured the density of neurons across the entire macaque cortex using the isotropic fractionator (a.k.a. brain soup) method (Collins et al. 2010). After mapping the neuron density data from the study by Collins et al. 2010 and the receptor data to a common atlas, we divided the receptor density by the neuron density, to obtain an estimate of D1 receptors per neuron in each cortical area.

### Retrograde tract-tracing

The inter-areal connectivity data in this paper is part of an ongoing effort to map the cortical connectome of the macaque using retrograde tract-tracing (Markov et al. 2013, 2014a,b). For each target area, a retrograde tracer was injected into the cortex. The tracer was taken up in the axon terminals in this area, and retrogradely transported to the cell bodies of neurons that projected to the target. These cell bodies could be throughout the brain. Each of these cell bodies in cortex was counted as a labelled neuron (LN). The amount of labelled neurons was counted in all cortical areas except for the injected target area. The cortical areas that send axons to the target area are called source areas. As there are uncontrollable differences in tracer volume and uptake between injections, we estimated the strength of connections as follows. For a given injection, the total number of cell bodies in the cortex outside of the injected (target) area was counted. The number of labeled neurons was within a source cortical area was then divided by the number of labeled neurons in the whole cortex (excluding the target area), to give a fraction of labeled neurons (FLN). The FLN was averaged across all injections in a given target area. For this calculation, we include all areas in the entire cortical hemisphere (*n*^*areas*^ = 91).

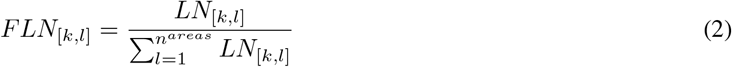

In addition, for each connection we defined the supragranular labeled neurons (SLN) as the fraction of neurons in the source area whose cell bodies were in the superficial (aka supragranular) layers.

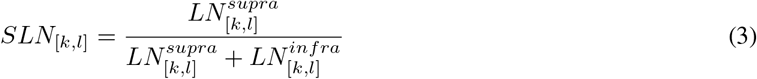

The subiculum (SUB) and piriform cortex (PIR) have a qualitatively different laminar structure to the neocortical areas, and thus supra- and infra-laminar connections (and thus the SLN) from these areas are undefined. We thus removed all connections from these areas from the following calculations (*n*^*areas,SLN*^ = 89). These connectivity matrices will be available on core-nets.org.

### Estimation of the cortical hierarchy

Following (Markov et al. 2014a), we estimate the hierarchical position *h* of each area using the SLN values of its connections. Feedforward connections tend to originate in the supragranular layers, while feedback connections tend to originate in the deep layers of the source area (Barone et al. 2000; Felleman and Van Essen 1991). Moreover, if a target area occupies a much higher hierarchical position than the source area, a greater proportion of the neurons emerge from the supragranular layers of the source area than if the two areas are closer in the hierarchy (Barone et al. 2000). Likewise for the feedback connections, a greater hierarchical distance between the areas implies that the higher area sends a greater proportion of it projections from the infragranular layers. This implies that the fraction of neurons coming from the supragranular layers in a given connection gives an estimate of the relative hierarchical position of two connected areas (Barone et al. 2000; Markov et al. 2014a).

Here, following (Markov et al. 2014a), we estimate a set of hierarchical levels (one per area) that best predicts the SLN values for all connections in the dataset.

The model to estimate the hierarchy has the form

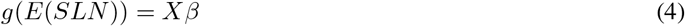

where g is a function that links the SLN of the connection between areas to the hierarchical distance between them. *β* is a column vector of length *n*^*areas,SLN*^, containing the hierarchy values to be estimated. *X* is an incidence matrix of shape *n*^*conns*^ × *n*^*areas,SLN*^, where *n*^*conns*^ (= 2619) is the number of connections between cortical areas in the remaining dataset. Each row in *X* represents a connection, and each column represents a cortical area. All entries in each row equal 0 except for the column corresponding to the source area, which has a value of −1, and the target (recipient) area, which has a value of 1 (Strang 1993).

The hierarchical values can be estimated with maximum likelihood regression. However, the model is singular (the rows sum to zero). In order to make the model identifiable, we therefore removed one column from *X*. We chose to remove the column corresponding to area V1, which is therefore forced to have a hierarchical value of 0. However, the choice of column is unimportant, as it is possible to estimate negative hierarchical values (in the case that other areas are lower than V1 in the hierarchy).

We used the beta-binomial model. The binomial parameter *p* corresponds to the proportion of successes. This is thought to be a random variable following a Beta distribution. The beta-binomial distribution depends on two parameters, the mean (*μ*, here the SLN), and the dispersion (*ϕ*). The beta-binomial model can account for the overdispersion of the neural count data. Note that the SLN of each measured connection is input into the model, without averaging across repeated injections.

The likelihood is written as

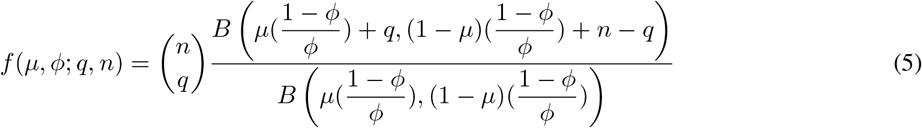

where *q* is the number of neurons projecting from the supragranular layers, n is the number of neurons projecting from all layers, and B is the beta function defined as

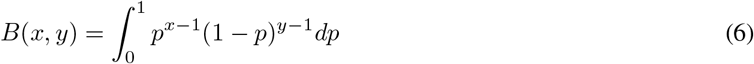

with *x, y >* 0. We fit the model using *μ* = Φ(*Xβ*), where Φ is the cumulative Gaussian, as it maps the real numbers to the (0,1) range. Φ^−1^ = *g* in equation 3 is the probit link function. The hierarchy is estimated by minimising the log-likelihood. For more details see (Markov et al. 2014a).

We then rescaled the hierarchy so that the maximum hierarchial value within the 40 region complete subgraph (containing all injected areas) equaled 1:

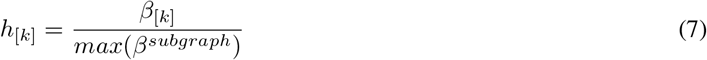

for all cortical areas *k* in the complete 40-area subgraph.

### Integration of anatomical datasets

All anatomical data was mapped to the appropriate parcellations on the Yerkes19 surface. For the present study, we mapped all data to the 40 area Lyon subgraph Markov et al. 2014b, as the areas in this parcellation were generally larger than those in the Jülich macaque atlas (Impieri et al., 2019; Niu et al., 2020; Rapan et al., In Prep; Niu et al., In prep; this paper) and the Queensland (spine count) injection sites (Elston 2007), and closer to standard areal descriptions than the Vanderbilt (neuronal density) (Collins et al. 2010) sections.

The receptor densities were quantified in 109 cortical regions defined by cyto- and receptor-architecture. The delineation of cortical region borders in the superior parietal lobe is described in (Impieri et al. 2019). Using the same method, anatomists (NPG, MN, LR) identified cortical areas on the basis of the receptor and cyto-architecture. See Figure 1 and associated data for the definition of the areas. Anatomists (NPG, MN, LR) carefully drew and independently revised defined borders on the Yerkes19 cortical surface (Donahue et al. 2016) to enable comparison with other data types. The D1 receptor data was mapped to the Lyon atlas as follows. For each area in the Lyon atlas, we searched for overlaps with areas in the Jülich macaque atlas. If more than 50% of the vertices within the area were also in the Jülich macaque atlas, the D1 receptor density for the area was calculated. All vertices within each Jülich area were assigned the mean value for that area. We averaged the D1 receptor density across all vertices that lay within both the Lyon area and the Jülich macaque atlas, thus performing a weighted average of the D1 receptor densities according to the degree of spatial overlap. Thirty-two of the 40 Lyon areas were assigned D1 receptor density in this way, with the remaining eight areas not overlapping sufficiently with the Jülich macaque atlas. Due to the strong positive correlation between the D1 receptor/neuron density and the hierarchy (Fig 1), for the simulations we inferred values for the remaining eight regions using linear regression with hierarchy as the independent variable and D1 receptor/neuron density as the independent variable.

Neuron density data was taken from (Collins et al. 2010). In the original paper, the cortex was divided into 42 regions and displayed on a flatmap, with anatomical landmarks labeled (Fig 2 and S1 of that paper). The borders of these regions were drawn on the Yerkes19 surface by SFW with reference to the original paper (Collins et al. 2010), several anatomical papers from the same group (Beck and Kaas 1999; Cerkevich et al. 2014; Kaas 2004), the Jülich (109 areas) and the Lyon (Markov-132) atlases (Donahue et al. 2016; Markov et al. 2014b), and were independently assessed by anatomists (LR, MN, NPG). The neural density data covered the entire cortex. As such, we assigned neural density to each area in the Lyon atlas, weighted by the spatial overlap with the original areas in the Vanderbilt atlas. D1 receptor density was divided by the neuron density to give the D1 receptor/neuron density in each area.

The Lyon atlas used to define the interareal connectivity data (Markov et al. 2014b) is already available on the Yerkes19 surface (Donahue et al. 2016). The complete subgraph including bidirectional connectivity has since been expanded from 29 areas in Donahue et al. 2016 to 40 areas in this paper.

For the spine count data, outlines of the 27 injection sites were drawn on the Yerkes19 surface by SFW with reference to the original papers (most of which had substantial anatomical description and hand-drawn maps), as well as anatomical papers cited within the original papers (Cavada and Goldman-Rakic 1989; Preuss and Goldman-Rakic 1991) and the Lyon and Jülich macaque atlases. Again, boundaries were independently assessed by anatomists (LR, MN, NPG). Spine count data was expressed according to injection sites, rather than entire cortical areas. As such, we found the number of vertices from each injection site overlapping with each area in the Lyon atlas. For each Lyon area, the spine count was an average of the spine counts for all the injection sites overlapping with the area, weighted by the number of vertices of each injection site contained within the area. In this way we estimated the spine counts on pyramidal cells in 24 of the 40 regions in the Lyon atlas. Based on the strong positive correlation between spine count and cortical hierarchy (r = 0.61, p = 0.001), and following previous work (Chaudhuri et al. 2015; Mejias and Wang 2019), we inferred the spine count for the remaining regions based on the hierarchy using linear regression.

Delineations of the areal borders for each atlas, and the anatomical data in the Yerkes19 space will be made available on the BALSA database upon publication.

### Overview of dynamical models

We first describe the connectivity structure of our local circuit model, and how dopamine modulates the efficacy of these connections. We then describe a large-scale dynamical model, in which the local circuit is used as a building block, and placed in each of 40 cortical areas. We describe the various steps to building the large-scale model, including how to connect the cortical areas, apply heterogeneity of excitation and the gradient of dopamine. Lastly, we describe how we simulated working memory tasks, lesions and optogenetic inhibition in this model.

### Description of the local cortical circuit

We describe a local cortical circuit containing populations of four distinct types of neurons. This is conceptually related to previous computational models of working memory involving multiple types of interneurons (Tanaka 1999; Wang et al. 2004a), and uses a mean field reduction of a spiking model (Brunel and Wang 2001; Wong and Wang 2006). PV, SST/CB and VIP/CR cells differed in the threshold and slope of their input-output function (f-I curve) (Bacci et al. 2003), local (Adesnik et al. 2012; Jiang et al. 2015; Muñoz et al. 2017; Pfeffer et al. 2013; Tremblay et al. 2016) and long-range connectivity (Lee et al. 2013; Wall et al. 2016), adaptation rates (Kawaguchi 1993; Mendonça et al. 2016; Schuman et al. 2019), and NMDA/AMPA ratio (Lu et al. 2007).

The connectivity structure and strengths of the local circuit, are based on a synthesis of anatomical and physiological studies, and are captured in the local connectivity matrix *G* (Tables 1 and 2) (Jiang et al. 2015; Kalisman et al. 2005; Lee et al. 2013; Ma et al. 2012; Markram et al. 1997; Pfeffer et al. 2013; Silberberg and Markram 2007; Walker et al. 2016).

**Table 1.**
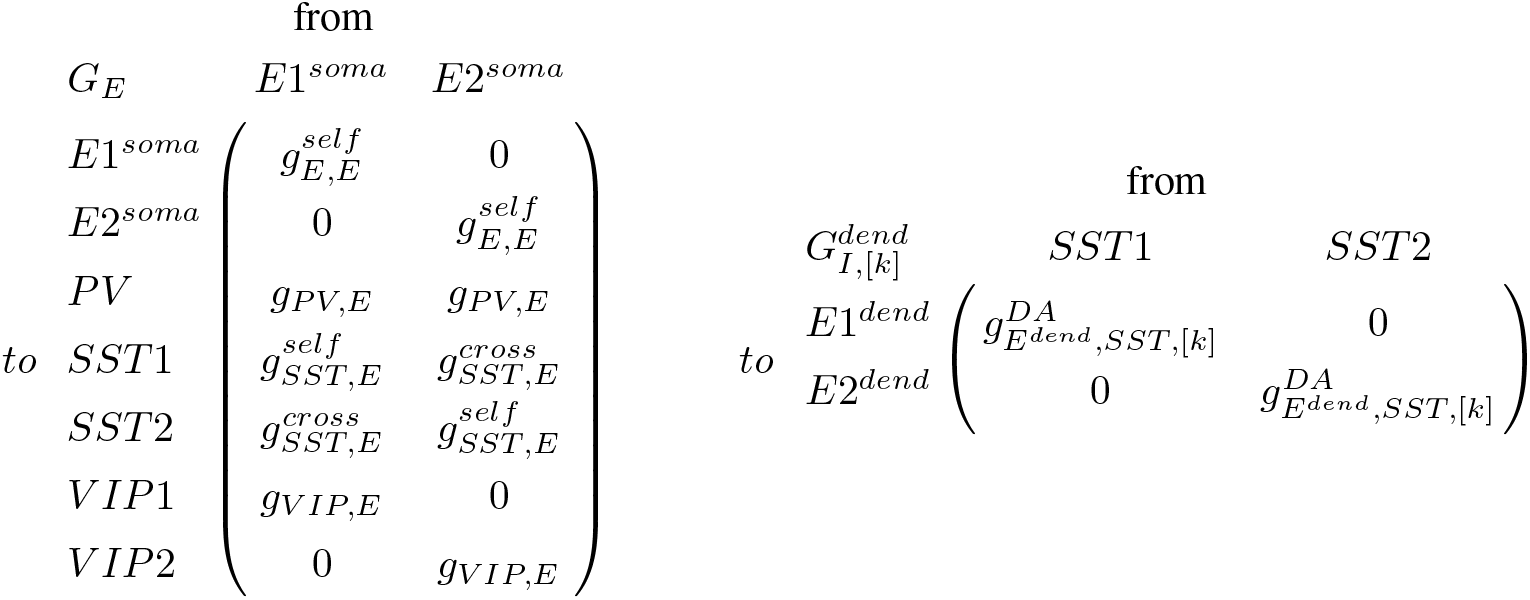
Left: Local excitatory output connections target excitatory and inhibitory populations. Right: SST/CB interneurons target the dendrites of pyramidal cells.

**Table 2.**
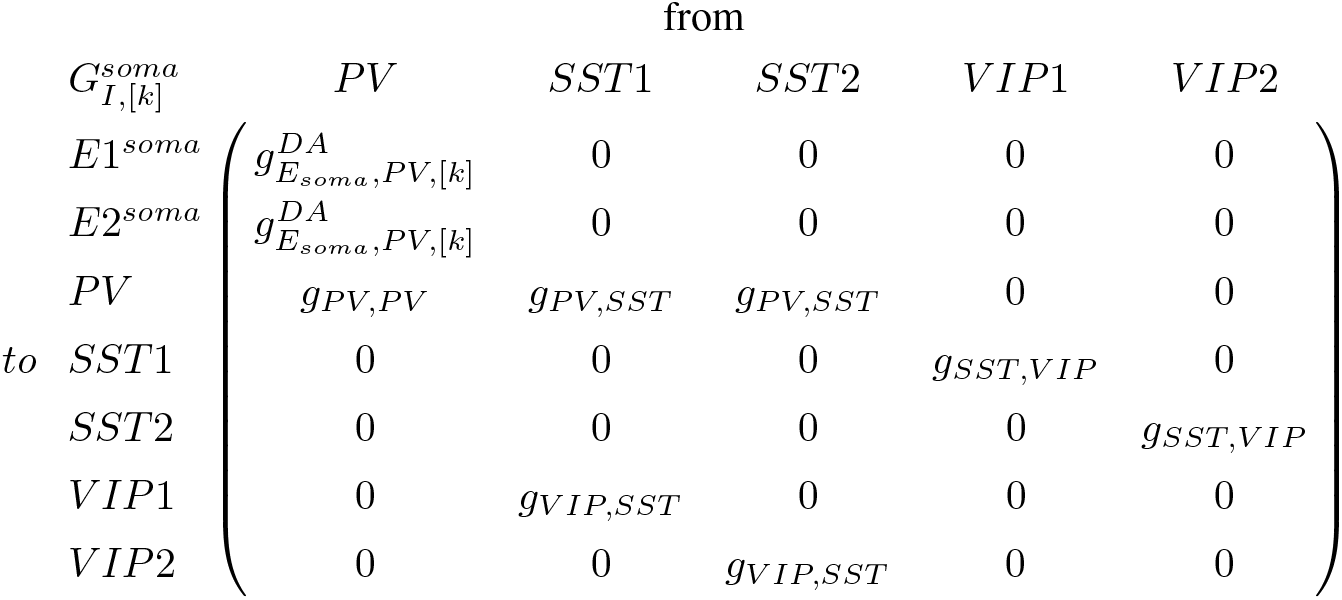
PV cells inhibit the cell body of pyramidal cells, but are themselves inhibited by other PV cells and SST/CB cells. SST/CB cells and VIP/CR cells mutually inhibit each other.

Note that connection probability and synaptic strength between neural types are generally positive correlated (Jiang et al. 2015). This simplifies the process of identifying the relative strengths of connections between neural populations in the circuit.

See Table 4 for all parameter values.

### Dopamine modulation

The density of dopamine D1 receptors per neuron was rescaled, so that the area with minimum density 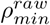 was set to zero, and the area with maximum density 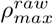 was set to one, with all other areas lying in between.

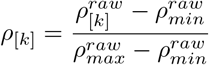

for all cortical areas *k*.

Network behavior was investigated for differing amounts of cortical dopamine availability (*λ*^*DA*^). The specific value of *λ*^*DA*^ used for each simulation is shown in the figures and main text. Note that for Figure 7, *λ*^*DA*^ is calculated dynamically throughout each trial. Cortical dopamine availability is related to the fraction of occupied D1 receptors *λ*^*occ*^ through a sigmoid function. The fraction of occupied D1 receptors thus lies between 0 and 1, as expected.

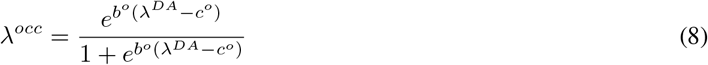

Dopamine increases the proportion of inhibition onto the dendrites of pyramidal cells (Gao et al. 2003). Therefore, we simulated the effect of dopamine on dendritic inhibition as follows. The total amount of dendritic inhibition increases (from a minimum to a maximum strength) as the total amount of occupied receptors increases. The total amount of occupied receptors is equal to the receptor density multiplied by the fraction of occupied receptors.

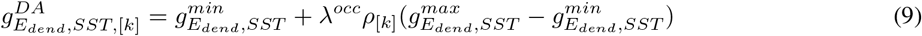

Dopamine decreases the proportion of inhibition onto the soma of pyramidal cells (Gao et al. 2003). Therefore, we simulated the effect of dopamine on somatic inhibition as follows. The total amount of somatic inhibition decreases (from a maximum to a minimum strength) as the total amount of occupied receptors increases.

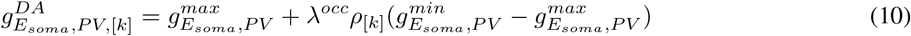

Dopamine also increases the strength of excitatory synaptic transmission via NMDA receptors (Seamans et al. 2001a). We modeled this with a sigmoid function, so that dopamine primarily increases NMDA conductances at low and medium dopamine concentrations, before reaching a plateau (Brunel and Wang 2001).

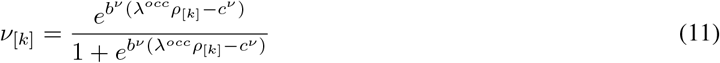

Here *b*^*ν*^ sets the slope of the sigmoid function, *c*^*ν*^ sets the midpoint.

The effects of dopamine on NMDA transmission is then defined as

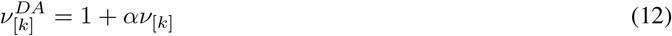

where *α* controls the strength of dopamine modulation on NMDA transmission.

High levels of D1 agonism lead to a reduction in pyramidal cell firing, particularly during the delay period of working memory tasks. D1 receptor stimulation may lead to inhibition of ongoing activity by engaging an intracellular pathway involving cyclic AMP, protein kinase A and either HCN or KCNQ channels (Arnsten et al. 2019; Gamo et al. 2015; Vijayraghavan et al. 2007). The mechanisms by which HCN channels may hyperpolarise the cell are still under debate (George et al. 2009; Pereira 2014). We simulated an increase in adaptation for very high levels of D1 receptor stimulation with a sigmoid function, so that adaptation increases at high dopamine concentrations, before reaching a plateau.

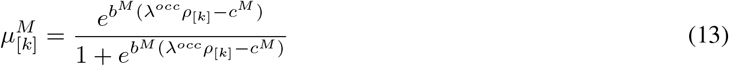

### Description of dynamical variables

The neural populations interact via synapses that contain NMDA, AMPA and GABA receptors. Each receptor has its own dynamics, governed by the following equations.

The synaptic variables are updated as follows (Wang 1999; Wong and Wang 2006; Yang et al. 2016)

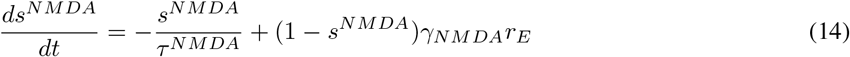

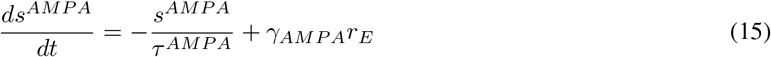

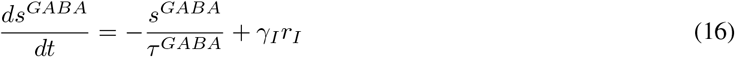

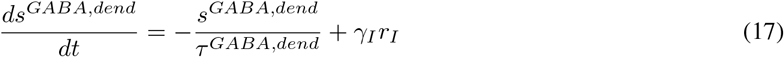

where *s* is the synaptic drive onto a particular receptor type, *τ* is the time constant of decay of that receptor and *γ*_*NMDA*_, *γ*_*AMPA*_ and *γ*_*I*_ are constants. *r*_*E*_ and *r*_*I*_ are the firing rates of the presynaptic excitatory and inhibitory cells targeting the NMDA, AMPA and GABA receptors, calculated below. Note that the inhibition onto the dendrite is slower than inhibition elsewhere (*τ*^*GABA,dend*^ > *τ*^*GABA*^) (Ali and Thomson 2008). Hence we calculate dynamics of dendritic and somatic inhibition separately.

Adaptation acts to reduce the firing rate when the rate is high, using the equation described in (Engel and Wang 2011),

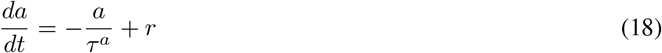

where *a* is the adaptation variable, *τ*^*A*^ is the adaptation time constant, and *r* is the firing rate of the neural population.

### NMDA/AMPA ratio

The fraction of excitatory postsynaptic current that is dependent on NMDA vs AMPA receptors differs by cell type (e.g. with relatively more current via the NMDA receptors in SST/CB vs PV cells) (Lu et al. 2007). Thus, we allowed the strength of excitatory transmission via NMDA and AMPA receptors to vary by cell type, described in the NMDA fraction, *κ* (Table 4).

#### Modulation of excitatory connections by dendritic spines

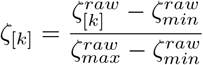

for all cortical areas [*k*].

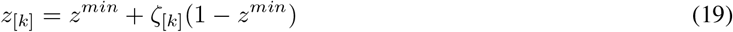

where *z*^*min*^ sets the lower bound for the modulation of excitatory connections by the spine count, *ζ*.

#### Description of local currents

The local NMDA current is calculated as follows

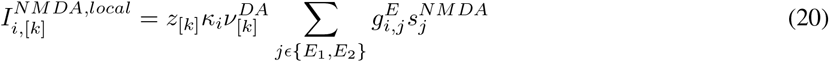

where the local excitatory connections via the NMDA receptors are scaled by the NMDA receptor fraction *κ*_*i*_, the dendritic spine count *z*_[*k*]_ and the D1 receptor stimulation 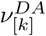 for all populations of neurons *i* and cortical areas *k*.

Similarly local excitatory connections via the AMPA receptors are scaled by the AMPA receptor fraction 1 − *κ*_*i*_ and the dendritic spine count *z*_[*k*]_.

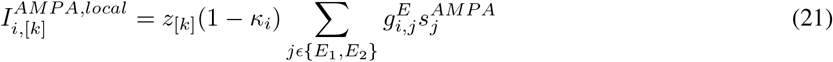

Local inhibitory connections are not explicitly modulated by the dendritic spine count (as spines are the locations of synapses between excitatory cortical neurons). Note however, that the connectivity structure *g*_*GABA*_ is modulated by the dopamine receptor density and occupancy (See Tables 1, 2 and 4).

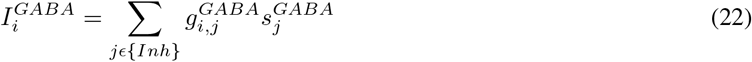

where *Inh* is the set of inhibitory neuron populations.

The currents onto the dendrites are calculated separately, in order to calculate the nonlinear transformation of the current in the dendrite. They depend on the noise and background currents, so are described below.

#### Description of noise and background currents

Noise is modeled as an Ornstein-Uhlenbeck process, separately for each population.

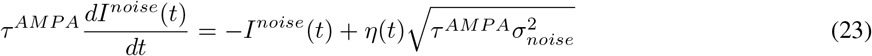

where *σ*_*noise*_ is the standard deviation of the noise and *η* is Gaussian white noise with zero mean and unit variance.

A constant background current *I*^*bg*^ was also added to each population (Table 4). This represents input from brain areas that are not explicitly modeled.

#### Description of the adaptation current

We include adaptation in excitatory cells (Kawaguchi 1993), SST/CB (Kawaguchi 1993, 1995) and VIP/CR cells (Mendonça et al. 2016; Schuman et al. 2019), but not PV cells (Kawaguchi 1993, 1995). This is reflected in their differing adaptation strengths 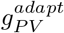 and *g*^*adapt*^, where 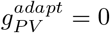.

The adaptation current is

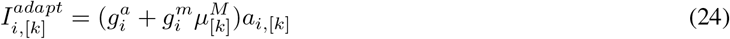

for all local populations *i* and cortical areas *k*.

Note that 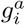 represents the non-dopamine dependent adaptation, while 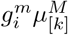 controls the dopamine-dependent adaptation, which depends on both dopamine release and receptor density (equation 13).

#### Large-scale connectivity structure

Each of the cortical areas is connected using connectivity strengths derived from the retrograde tract-tracing data. Parts of this dataset of been included in previous publications (Markov et al. 2013, 2014a,b). The long-range connectivity matrices are built from the FLN matrix. However, as noted in (Mejias et al. 2016), the FLN matrix spans 5 orders of magnitude. The relationship between anatomical and physiological connectivity strengths is not clear, but if we were to use the raw FLN values in the large-scale model, many of the weaker connections would become irrelevant. To deal with this, we rescale the FLN matrix in order to increase the influence of smaller connections while maintaining the topological structure (Mejias et al. 2016; Mejias and Wang 2019).

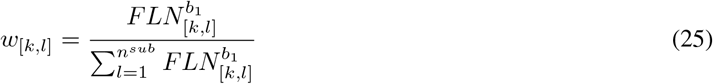

Here we restrict calculations to the injected cortical areas *i*, *j*, allowing for complete bidirectional connectivity within the subgraph (*n*^*sub*^ = 40). We use the same parameter values as in (Mejias et al. 2016; Mejias and Wang 2019) (Table 4) to construct our interareal connectivity matrix *W*.

As noted previously, feedforward projections tend to originate in the supragranular layers, while feedback connections originate in the deep layers. Feedforward and feedback connections also likely have different cellular targets. Therefore it is useful to separate the long-distance feedforward and feedback connections.

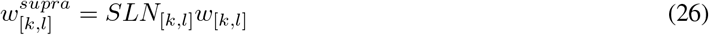

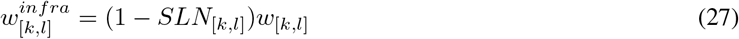

#### Interareal population interactions

The majority of interareal connections contain a mixture of axons projecting from deep and superficial layers. Long distance connections onto excitatory cells primarily target the distal dendrites (Petreanu et al. 2009) (Table 3). Therefore, in the model we assume that long-distance connections target the dendrites of excitatory cells. VIP/CR cells receive the strongest long-distance inputs of all inhibitory cells, while SST/CB receives the weakest (Lee et al. 2013; Wall et al. 2016) (Table 3, Table 4). This suggests that long-range connections effectively disinhibit the dendrite in the target area by exciting VIP/CR interneurons, while concurrently exciting the dendrite, to maximize the probability of information passing from the source area into the target area. Following Mejias and Wang 2019 we assume that feedback connections target inhibitory cells more strongly than feedforward connections.

**Table 3.** Long-range targets onto excitatory (left) and inhibitory (right) cells

|  |  | from |  |  |  | from |  |
| --- | --- | --- | --- | --- | --- | --- | --- |
| $J^{E,E}$ | | $E1_{soma}$ | $E2_{soma}$ | $J^{I,E}$ | | $E1_{soma}$ | $E2_{soma}$ |
| to | $E1_{soma}$ | 0 | 0 | to | PV | $g_{PV,E}^{LR}$ | $g_{PV,E}^{LR}$ |
| | $E2_{soma}$ | 0 | 0 | | SST1 | $g_{SST,E}^{LR}$ | 0 |
| | $E1_{dend}$ | $g_{E,E}^{LR,self}$ | $g_{E,E}^{LR,cross}$ | | SST2 | 0 | $g_{SST,E}^{LR}$ |
| | $E2_{dend}$ | $g_{E,E}^{LR,cross}$ | $g_{E,E}^{LR,self}$ | | VIP1 | $g_{VIP,E}$ | 0 |
| | | | | | VIP2 | 0 | $g_{VIP,E}$ |

**Table 4.**
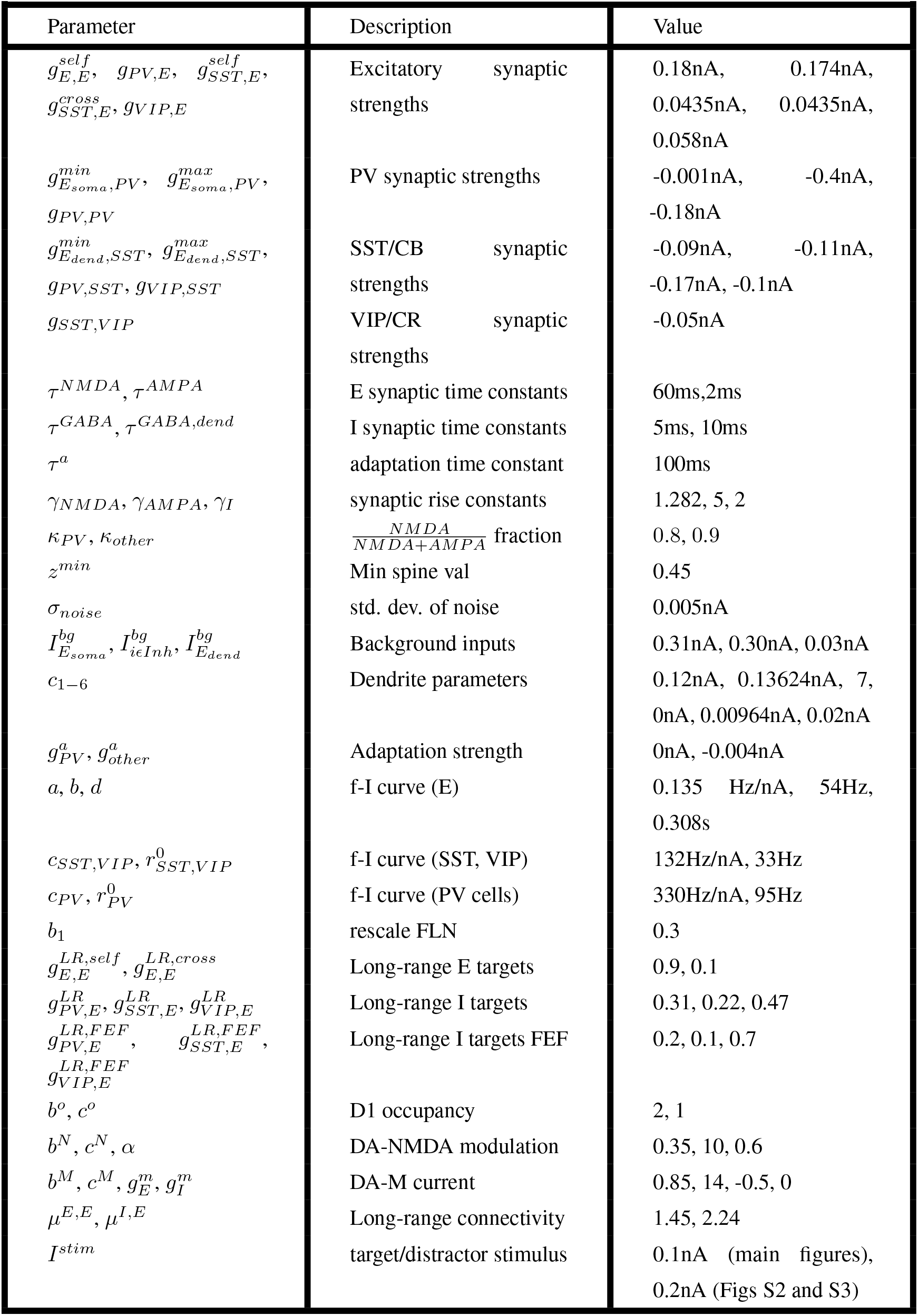
Parameters for Numerical Simulations

| Parameter | Description | Value |
| --- | --- | --- |
| $g_{E,E}^{self}, g_{PV,E}, g_{SST,E}^{self}, g_{SST,E}^{cross}, g_{VIP,E}$ | Excitatory synaptic strengths | 0.18nA, 0.174nA, 0.0435nA, 0.0435nA, 0.058nA |
| $g_{E_{soma},PV}^{min}, g_{E_{soma},PV}^{max}, g_{PV,PV}$ | PV synaptic strengths | -0.001nA, -0.4nA, -0.18nA |
| $g_{E_{dend},SST}^{min}, g_{E_{dend},SST}^{max}, g_{PV,SST}, g_{VIP,SST}$ | SST/CB synaptic strengths | -0.09nA, -0.11nA, -0.17nA, -0.1nA |
| $g_{SST,VIP}$ | VIP/CR synaptic strengths | -0.05nA |
| $\tau^{NMDA}, \tau^{AMPA}$ | E synaptic time constants | 60ms, 2ms |
| $\tau^{GABA}, \tau^{GABA,dend}$ | I synaptic time constants | 5ms, 10ms |
| $\tau^a$ | adaptation time constant | 100ms |
| $\gamma^{NMDA}, \gamma^{AMPA}, \gamma^I$ | synaptic rise constants | 1.282, 5, 2 |
| $\kappa_{PV}, \kappa_{other}$ | $\frac{NMDA}{NMDA+AMPA}$ fraction | 0.8, 0.9 |
| $z^{min}$ | Min spine val | 0.45 |
| $\sigma_{noise}$ | std. dev. of noise | 0.005nA |
| $I_{E_{soma}}^{bg}, I_{i \in Inh}^{bg}, I_{E_{dend}}^{bg}$ | Background inputs | 0.31nA, 0.30nA, 0.03nA |
| $c_{1-6}$ | Dendrite parameters | 0.12nA, 0.13624nA, 7, 0nA, 0.00964nA, 0.02nA |
| $g_{PV}^a, g_{other}^a$ | Adaptation strength | 0nA, -0.004nA |
| $a, b, d$ | f-I curve (E) | 0.135 Hz/nA, 54Hz, 0.308s |
| $c_{SST,VIP}, r_{SST,VIP}^0$ | f-I curve (SST, VIP) | 132Hz/nA, 33Hz |
| $c_{PV}, r_{PV}^0$ | f-I curve (PV cells) | 330Hz/nA, 95Hz |
| $b_1$ | rescale FLN | 0.3 |
| $g_{E,E}^{LR,self}, g_{E,E}^{LR,cross}$ | Long-range E targets | 0.9, 0.1 |
| $g_{PV,E}^{LR}, g_{SST,E}^{LR}, g_{VIP,E}^{LR}$ | Long-range I targets | 0.31, 0.22, 0.47 |
| $g_{PV,E}^{LR,FEF}, g_{SST,E}^{LR,FEF}, g_{VIP,E}^{LR,FEF}$ | Long-range I targets FEF | 0.2, 0.1, 0.7 |
| $b^o, c^o$ | D1 occupancy | 2, 1 |
| $b^N, c^N, \alpha$ | DA-NMDA modulation | 0.35, 10, 0.6 |
| $b^M, c^M, g_E^m, g_I^m$ | DA-M current | 0.85, 14, -0.5, 0 |
| $\mu^{E,E}, \mu^{I,E}$ | Long-range connectivity | 1.45, 2.24 |
| $I^{stim}$ | target/distractor stimulus | 0.1nA (main figures), 0.2nA (Figs S2 and S3) |

Excitatory cells in different cortical areas with the same receptive fields are more likely to be functionally connected (Zandvakili and Kohn 2015). This is reflected in our model as follows. In the source area, there are two excitatory populations, 1 and 2, each sensitive to a particular feature of a visual stimulus (such as a location in the visual field).

Likewise in the target area, there are two populations 1 and 2, sensitive to the same visual features. We assume that 90% of the output of population 1 in the source area goes to population 1 in the target area, and the remaining 10% to population 2. The converse is true for population 2 in the source area (it targets 10% population 1, 90% population 2; Table 3, Table 4).

#### Disinhibitory circuit in the frontal eye fields

The frontal eye fields (areas 8m and 8l in the model), have a very high percentage of calretinin neurons, and relatively fewer parvalbumin and calbindin neurons (Pouget et al. 2009). To account for this in the model, we relatively increased the long-range inputs to VIP/CR cells in areas 8m and 8l, as detailed in Table 4. These changes are critical for persistent activity in areas 8l and 8m, but otherwise do not greatly affect the behavior of the model. Without this change, the overlap between the simulated delay activity pattern and the experimental delay activity pattern (as in Figure 3) is still extremely high (17/19 areas correct, chi-square = 12.31 p = 0.0004), and the activity pattern depends on both the long-range connectivity (p = 0.001), and D1 receptor distribution (p = 0.008), but not the spine count (p = 0.19). All other results are unchanged.

#### Calculation of long-range currents

Long-range interactions are applied as follows:

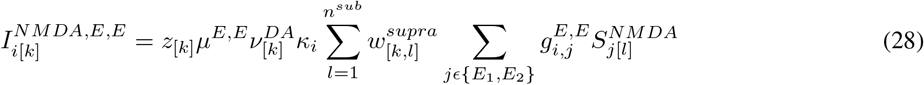

where *z*_[*k*]_ is the dendritic spine count for area *k* (as defined above), *μ*^*E,E*^ is the long-range connectivity strength onto excitatory cells (See Table 4), 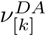 is the degree of dopamine modulation of NMDA currents for area *k*, *κ*_*i*_ is the NMDA/AMPA fraction for population *i*, *w*_[*k,l*]_ is the connection strength from area *l* to area *k*, 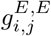 sets the long-range strength from population *j* to population *i* (Tables 3 and 4) and 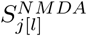 is the synaptic NMDA drive from population *j* in source area *l*.

Similarly,

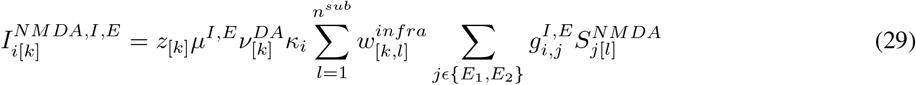

The total long-range current via the NMDA receptors, is simply the concatenation of the two above terms *I*^*NMDA,E,E*^ and *I*^*NMDA,I,E*^.

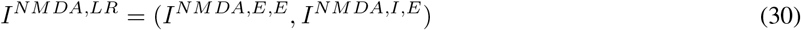

The long-range AMPA current is calculated similarly,

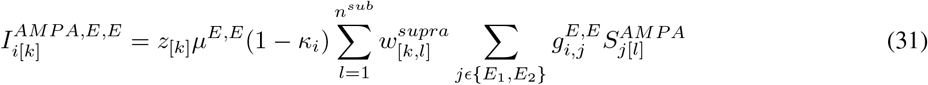

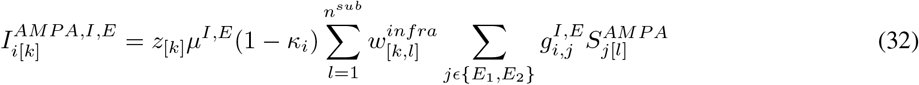

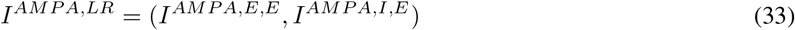

#### Description of dendritic currents

The inhibitory current onto the dendrite comes from SST/CB cells and is modulated by dopamine (Table 1, equation 8)

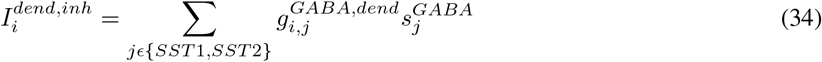

The distal dendrites receive long-range input (from neurons in other areas), noise and background input. In addition, if the area receives a stimulus directly, then the external stimulus also targets the dendrites. Note that most local connections target the area around the soma (Markram et al. 1997; Petreanu et al. 2009). This is reflected in the model by having local connections exclusively target the soma compartment of pyramidal cells.

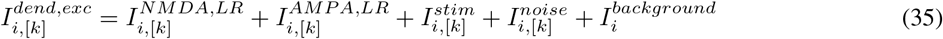

The dendritic nonlinearity is adapted from (Yang et al. 2016) and modeled as follows:

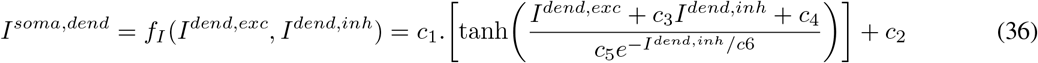

where *I*^*soma,dend*^ is the total current passed from the dendrite to the soma, *I*^*dend,exc*^ and *I*^*dend,inh*^ are the total excitatory and inhibitory current onto the dendrite, respectively. *c*1 to *c*6 control the gain, shift, inversion point and shape of the nonlinear function. These parameters are set to ensure that strong inhibition to the dendrite effectively blocks dendritic activity, but has little effect on somatic firing if the soma is directly stimulated (See Table 4) (Marlin and Carter 2014).

#### Application of external stimuli for tasks

In all simulations, the first stimulus is applied for 400ms. The second stimulus (Figures 5–7) is applied 600ms after the removal of the target stimulus for another 400ms. The two stimuli are of equal strength and duration, although the results are robust to a range of stimulus strengths (See Table 4 for parameter values). For Figures 2–7 in the main text, a stimulus was applied to the dendrite of excitatory population 1 in area V1. For Figures 5–7 a second stimulus was applied to the dendrite of excitatory population 2 of area V1. For Supplementary Figures 2 and 3, the stimuli were applied to area 3 of primary somatosensory cortex instead. In all equations, the target and distractor stimuli are designated by the term *I*^*stim*^.

#### Total current in large-scale model

The total current equals the sum of all long-range, local and external inputs, and intrinsic currents.

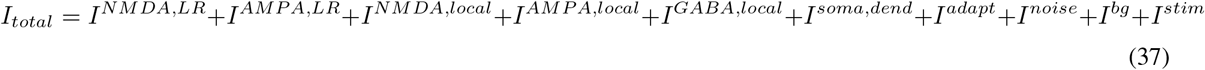

#### Description of f-I curves

The f-I (current to frequency) curve of the excitatory population is

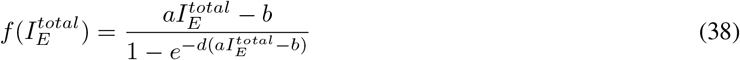

where *r*_*E*_ is the firing rate of an populations of excitatory cells, 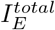 is the total input to the population, *a* is a gain factor, *d* determines the shape of 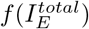, such that if *d* is large, 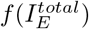 acts like a threshold-linear function, with threshold *b* (Abbott and Chance 2005).

The f-I curves for the inhibitory neuron populations are modeled using a threshold-linear function

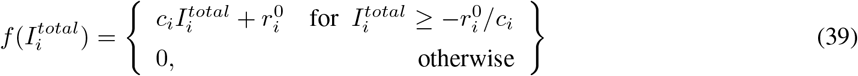

where *r*_*i*_ is the firing rate of a population of inhibitory cells, 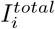 is the total input to the population.

The threshold 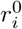 and slope *c*_*i*_ depend on the cell type *i* (Bacci et al. 2003). See Table 4 for parameter values.

The firing rates are updated as follows

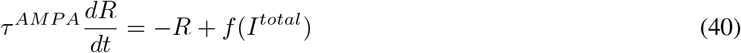

for all cell types.

#### Short-term synaptic plasticity

For Figure 5, we added short-term plasticity to synapses from excitatory cells to excitatory cells and SST/CB cells as follows (Mongillo et al. 2008).

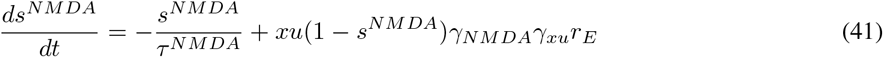

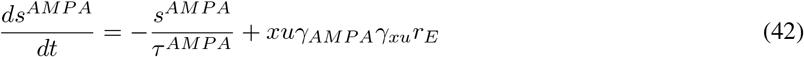

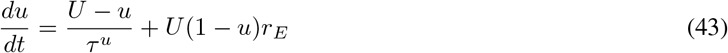

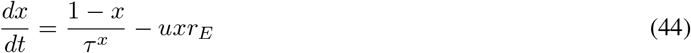

with *U* = 0.2, *τ*^*u*^ = 1.5*s*, *τ*^*x*^ = 0.2*s*, as in Mongillo et al. 2008. We also added a term *γ*_*xu*_ = 2.5 to account for the fact that the product *xu* is usually less then 1, and to keep firing rates similar to those in other simulations.

#### Comparing the simulated and experimental patterns of delay activity

In Figure 3, we compare the activity pattern of the model to the experimental pattern, and investigate its dependence on anatomical features. To shuffle anatomical connections, we shuffled connections within rows of the FLN matrix, so that the distribution of connections and connection strengths to each area remained constant, with the identity of the connections changing. The same reordering was applied to the SLN matrix. D1 receptor densities and spine counts were shuffled separately. Results were visualised using a custom version of a Raincloud Plot (Allen et al. 2019) to enable concurrent visualisation of the distribution and individual simulation results.

#### Lesioning of cortical areas

In Figure 4, we simulate the effects of a lesion to individual cortical areas. We do this by removing all input and output connections of the lesioned area in the connectivity matrices *W*^*E,E*^ and *W*^*I,E*^. For the statistical analysis of the relationship between anatomical features and visual effects, we removed areas V1 and V2 from the analysis. This was due to the fact that these areas were crucial to the propagation of the visual stimulus, but not working memory *per se* (Fig 4 and Fig S3). We performed a stepwise-linear regression approach. However, in order to allow for fair comparison between the anatomical predictors, for Fig 4D, we show the t-statistics for individual linear regression models with each anatomical predictor separately.

#### Simulated optogenetic inhibition of SST2 populations

In Figure 6, we simulate the effects of optogenetic inhibition to the SST2 populations in cortical areas in the frontoparietal network. The frontoparietal network is defined according to the results of Leavitt et al. 2017, as in Figure 3. To do this, we apply an external inhibitory stimulus of 0.1nA to these populations for the duration of the distractor stimulus. This may be possible experimentally (initially in mice), by identifying cells that are both active to the distractor stimulus (via a *cfos* immediate early gene promotor) and expressing SST, and optogenetically inhibiting them (Abbas et al. 2018; Liu et al. 2012).

#### Dynamics and connectivity within VTA

For Figure 7, we investigate whether the dynamics of dopamine release can be learned in order to selectively maintain the desired working memory content. Note both dopaminergic and GABAergic cells in the VTA receive excitatory input from the cortex, while the majority of inhibition to dopaminergic cells comes from local VTA GABAergic cells (Soden et al. 2020).

The total current input to the dopamine cells in VTA is

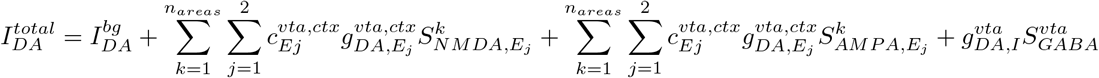

where 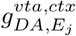 sets the maximum strength of cortical-VTA connections. 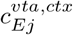 is the fraction of synapses in an up state (Soltani and Wang 2006), and is updated via reinforcement learning (see below). Initial values are 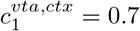, 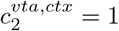. 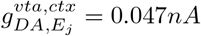 and 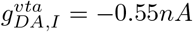.

The input to VTA inhibitory cells is

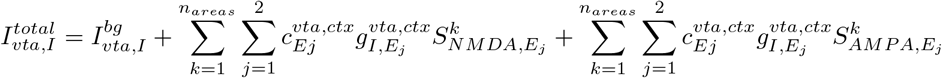

where 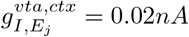

Synaptic inputs to the VTA inhibitory are driven by facilitating synapses (Soden et al. 2020), as in equations 41–44, but with *x* = 0.87 held constant and *τ*^*u*^ = 200*ms*

The firing rates of the dopamine cells *r*_*DA*_ as in equations 37 and 39. The firing rates of GABAergic cells are updated as in equations 38–39.

#### Cortical dopamine availability

For Figure 7, dopamine availability in the cortex *λ*^*DA*^ depends on the firing rates in the dopamine neurons as follows:

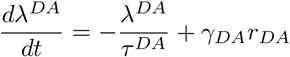

where *τ*^*DA*^ = 3*s* and *γ*_*DA*_ = 0.05.

#### Reward-based learning

The fraction of cortex to VTA synapses in the up state is updated according to the outcome of the previous trial, using the simplified learning rule of Soltani and Wang 2006

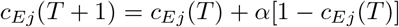

if target *j* is selected and rewarded and

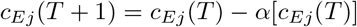

if target *j* is selected and not rewarded. *T* is the current trial and *α* = 0.2 is the learning rate.

## Acknowledgements

This project was funded by (NIH/BMBF) CRCNS grant (nos. R01MH122024 and 01GQ1902) to NPG and XJW; NIH grant R01MH062349, ONR grant N00014 and James Simons foundation grant 543057SPI to XJW and the European Union’s Horizon 2020 Framework Programme for Research and Innovation under the Specific Grant Agreements 785907 (Human Brain Project SGA2) and 945539 (Human Brain Project SGA3) to KZ and NPG. The authors would like to dedicate this manuscript to Prof. Karl Zilles, who was an inspiring mentor, colleague and friend. Prof. Zilles passed away earlier this year.

## Data and code availability

Upon acceptance for publication, we will make the autoradiography data available in table format. Additionally, we will publish the surface representation of all anatomical data used in the paper, and the corresponding cortical atlases to the BALSA neuroimaging website. All code used to produce the model and figures will be published on GitHub.

## Author contributions

Conceptualization - SFW, XJW. Methodology - SFW, DB, XD, KK, HK, KZ, NPG, XJW. Software - SFW. Validation - SFW, NPG, DB, XD, KZ, XJW. Formal Analysis - SFW. Investigation - NPG, LJR, MQ, HK, KZ. Resources - NPG, KK, HK, KZ, XJW. Writing - original draft preparation - SFW. Writing - review and editing - all authors. Visualization - SFW. Supervision - NPG, KZ, XJW. Funding acquisition - SFW, NPG, KK, HK, KZ, XJW.

## Supplementary Material

**Figure S1:**
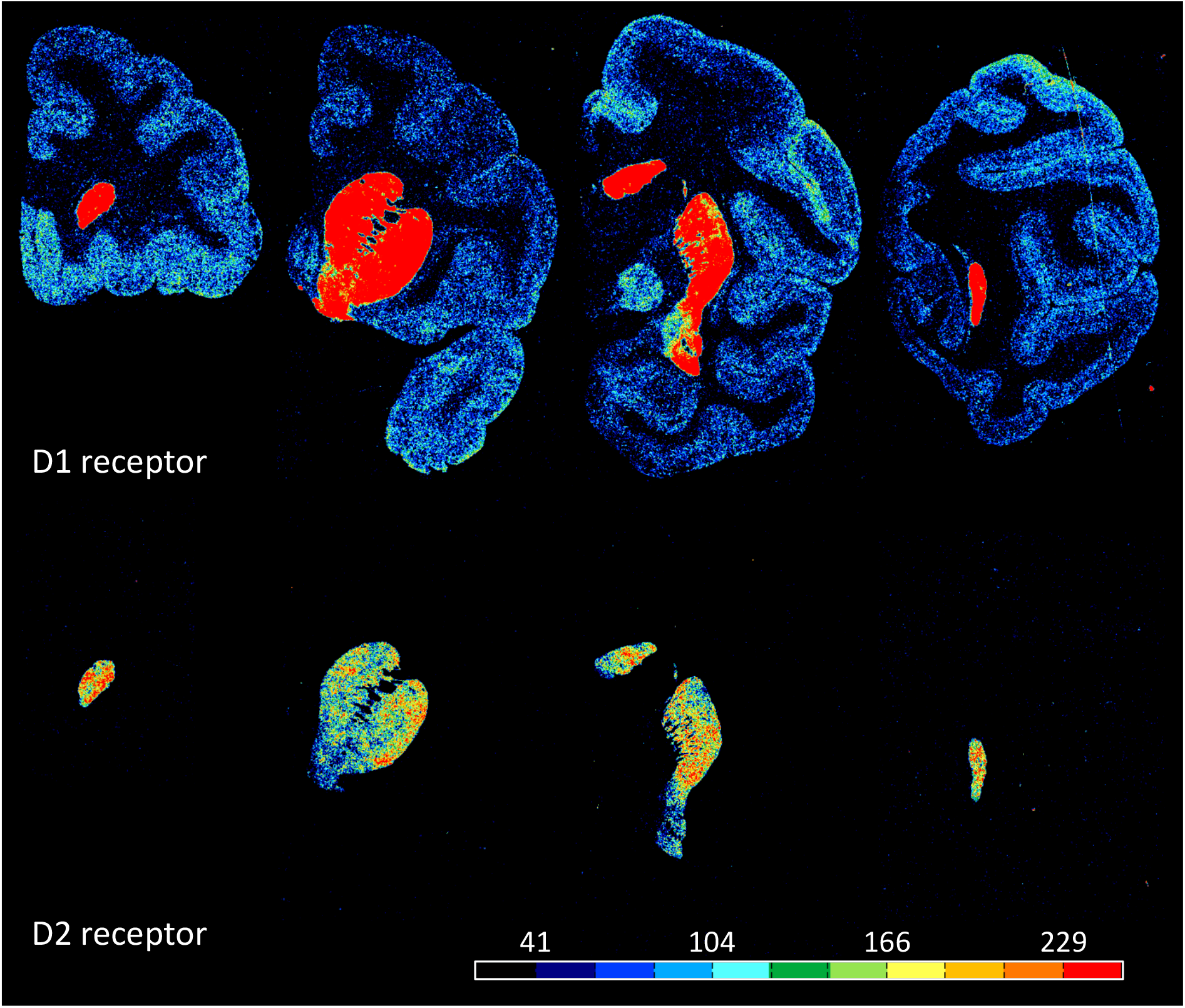
Exemplary coronal sections through the macaque brain and processed for visualization of dopamine D1 and D2 receptors by means of quantitative in-vitro receptor autoradiography. Note, that D2 receptor density in cortex is so low, that it is not detectable by means of the here applied method. Scale bar codes for receptor densities in fmol/mg protein.

**Figure S2:**
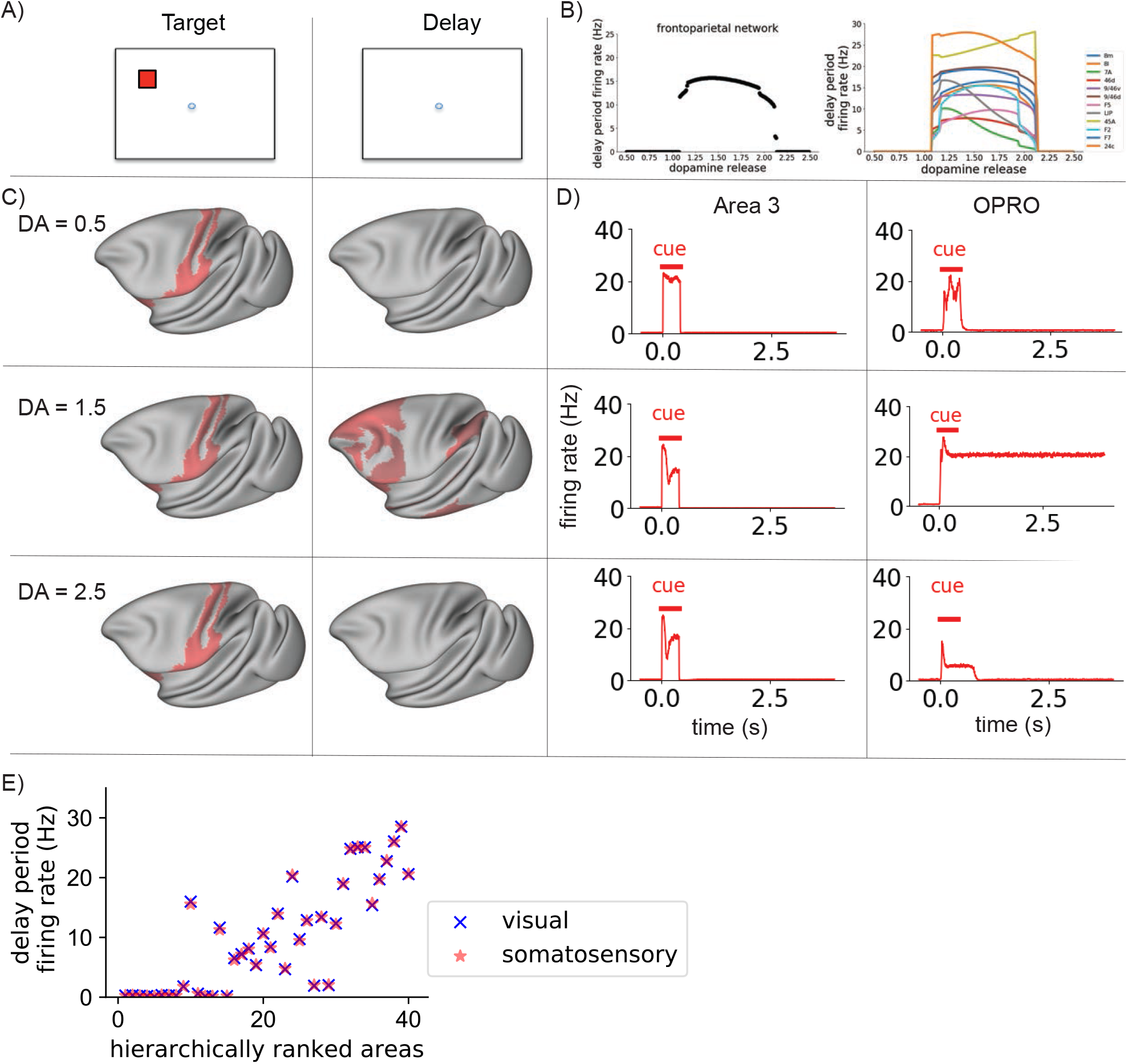
Dopamine release enables distributed somatosensory working memory. A) Structure of the task. The cortical network was presented with a stimulus, which it had to maintain through a delay period. The tactile stimulus is presented to primary somatosensory cortex (area 3). B, left) Mean firing rate in the frontoparietal network at the end of the delay period, for different levels of dopamine release. There is an inverted-U relationship between dopamine release and delay period activity across the frontoparietal network, as for visual working memory. B, right) Mean delay-period activity of cortical areas as a function of dopamine release. All areas shown display persistent activity in experiments (Leavitt et al. 2017). C) Activity is shown across the cortex at different stages in the working memory task (left to right), with increasing levels of dopamine release (from top to bottom). Red represents activity in the excitatory population sensitive to the location of the target stimulus. Very low or very high levels of dopamine release resulted in reduced propagation of stimulus-related activity to frontal areas and a failure to engage persistent activity. Mid-level dopamine release enables distributed persistent activity. D) Timecourses of activity in selected cortical areas. The horizontal bars indicate the timing of cue (red) input to area V1. Activity in early somatosensory areas such as area 3 peaks in response to the stimulus, but quickly decays away after stimulus removal for all levels of dopamine release. In contrast, there is dopamine-dependent persistent activity in area OPRO. E) The pattern of activity at the end of the delay period is highly overlapping following visual and somatosensory working memory tasks. DA, cortical dopamine availability.

**Figure S3:**
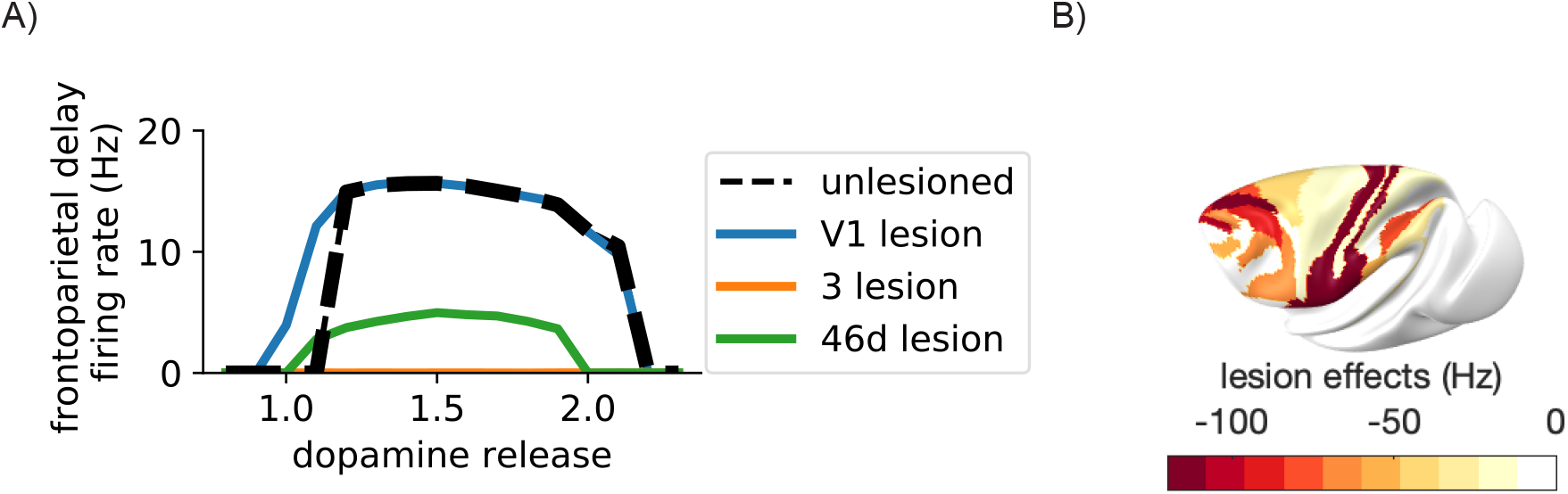
Lesions to visual areas do not disrupt somatosensory working memory. A) Lesions to areas such as 46d and LIP led to reduced delay period firing across for all levels of dopamine release. Lesions to ares 3 and 2 of somatosensory cortex disrupted the ability to perform the somatosensory working memory task. In contrast, lesions to visual areas such as V1 did not significantly affect somatosensory working memory. B) Map showing the severity of lesions to cortical areas on somatosensory working memory. More severe effects are shown in deeper red.

**Figure S4:**
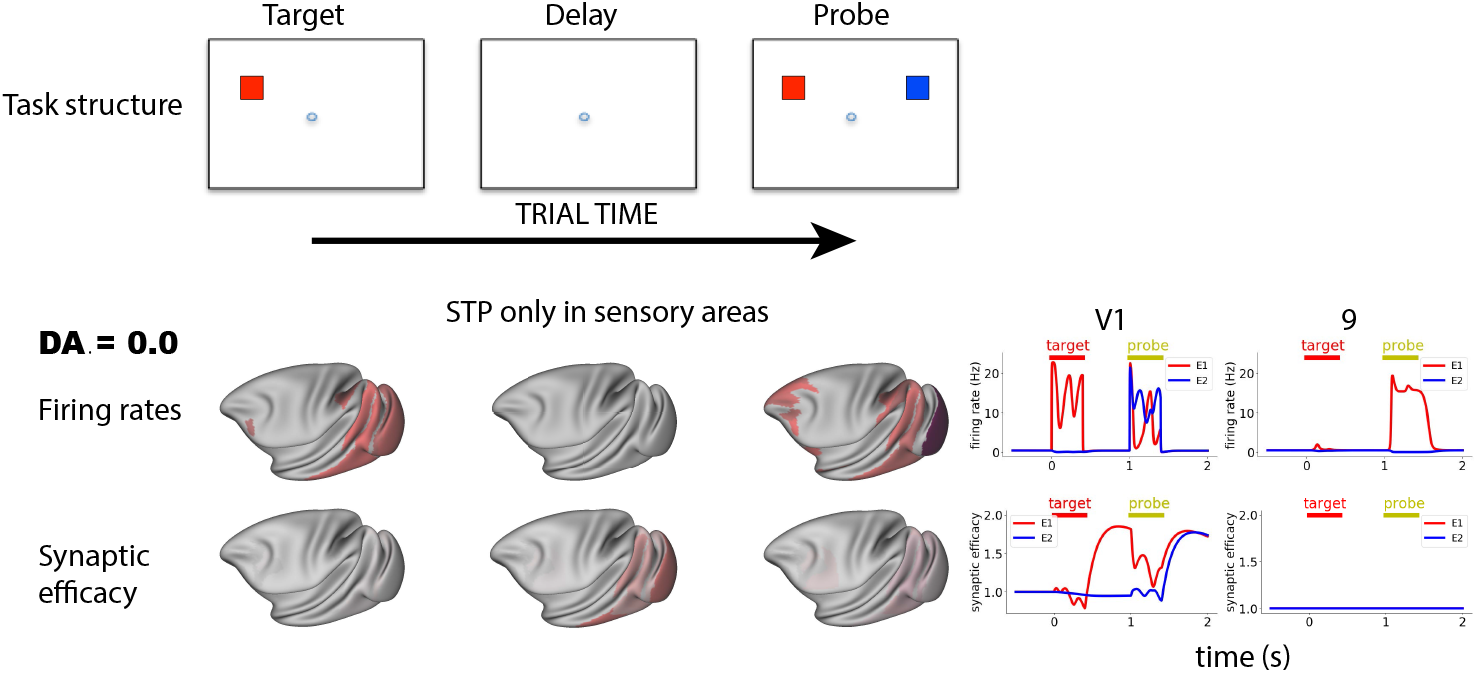
Activity-silent working memory without short-term plasticity in local prefrontal synapses. Reactivation of latent working memory representations was possible upon pinging the system, with short-term plasticity only on connections from neurons in sensory areas.

**Figure S5:**
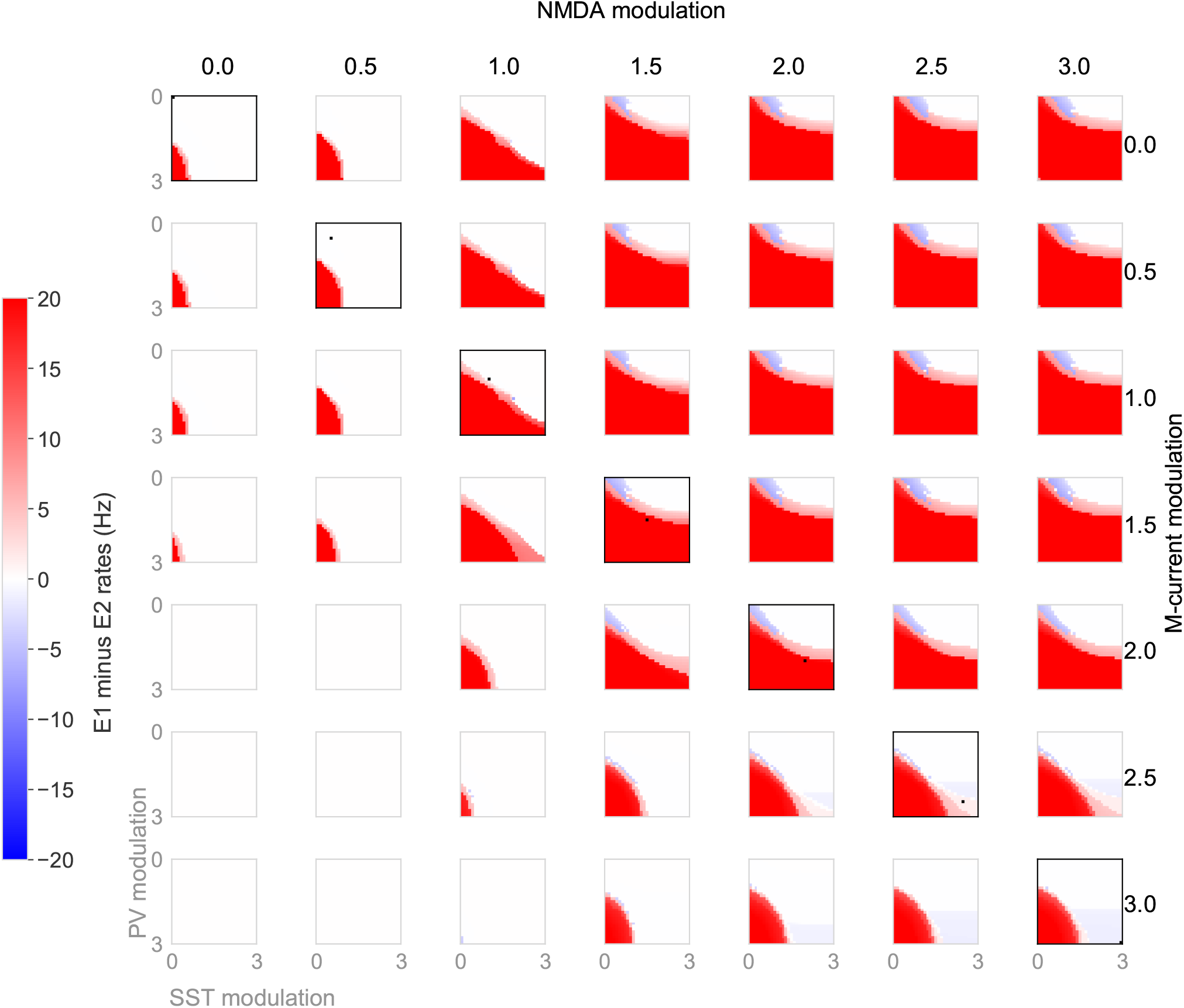
Distractor-resistance depends on the high dendritic inhibition. We identified the model behaviour for different dopamine levels, across different levels of dendritic and somatic inhibition. Consistently across dopamine levels, higher somatic, and lower dendritic inhibition was associated with distractible working memory (blue). In contrast, lower somatic, and higher somatic inhibition was associated with distractor-resistant working memory (red). High dendritic and high somatic inhibition results in no persistant activity (white). The levels of dendritic and somatic inhibition associated with the standard dopamine modulation used in the rest of the paper marked by a black square. Note that high PV modulation by dopamine results in lower PV inhibition of the soma.

## Notes

### Competing Interest Statement

The authors have declared no competing interest.

